# XRRA1 acts as a molecular brake on radiation-induced DNA damage signaling and immunogenic cell death in tumor cells

**DOI:** 10.64898/2026.04.21.719372

**Authors:** Tanvi Qamar, Saba Ubaid, Vipin Kumar, Mohammad Kashif, Tanvi Singh, Misba Majood, Ranjana Singh, Ajay Kumar Singh, Rashmi Kushwaha, Vivek Singh

**Author notes:** Correspondence to. Prof. Dr. Rashmi Kushwaha, Department of Pathology, King George’s Medical University, Lucknow, India. Email ID, Dr. Vivek Singh, Department of Biochemistry, King George’s Medical University, Lucknow, India. Email ID. Equally Contributed (Tanvi Qamar, Saba Ubaid, Vipin Kumar, Mohammad Kashif).

## Abstract

Radiotherapy kills cancer cells by inducing DNA damage, but adaptive responses that buffer injury and limit immunogenic signaling remain incompletely understood. Although ionizing radiation can activate cytosolic DNA sensing and immunogenic cell death, the tumor-intrinsic regulators linking these processes to radioresistance are not well defined. Here, we show that XRRA1 is a stress-adaptive determinant of radioresponse identified by integrating discovery proteomics into chronic myeloid leukemia with clinical tissue validation and functional studies across multiple tumor models. In peripheral-blood proteomes, XRRA1 was quantitatively reduced in chronic myeloid leukemia yet associated with a favorable prognosis and with networks enriched for RNA regulation, apoptosis, and DNA-repair biology. In human biopsy specimens, XRRA1 protein abundance was increased in radiation-exposed tissues, and in cultured cells, ionizing radiation induced XRRA1 more strongly and persistently in normal cells than in cancer cells. Silencing XRRA1 had little effect on basal growth but markedly enhanced radiation-induced loss of viability, apoptosis, spheroid disruption, and clonogenic failure, while increasing γH2AX- and DNA-PK-associated damage signaling and linking XRRA1 to non-homologous end joining factors. XRRA1 depletion also amplified cGAS-STING-TBK1-IRF3 activation, interferon-stimulated gene expression, extracellular ATP release, and cell-surface calreticulin exposure. These findings identify XRRA1 as a molecular brake that limits the conversion of radiation-induced DNA damage into immunogenic stress responses. XRRA1, therefore, represents a candidate biomarker of radioadaptive stress and a potential target for radiosensitization and radiotherapy-immunotherapy combinations.

## Introduction

Radiotherapy kills cancer cells largely by inducing DNA double-strand breaks, but intrinsic and acquired radioresistance continues to limit durable treatment responses. Experimental studies have shown that radiosensitivity is strongly influenced by the efficiency with which these lesions are detected and repaired: ATM-deficient cells display defective double-strand break repair and marked radiosensitivity, while core non-homologous end-joining (NHEJ) factors, including Ku, XRCC4, and DNA ligase IV, assemble at broken DNA ends to promote end alignment and ligation. Consistent with this, pharmacologic inhibition of DNA-PK enhances radiation-induced DNA damage, apoptosis, and loss of survival in tumor cells, supporting the view that efficient DNA repair is a major biological basis of radioresistance ^[1–4]^. Radiation response, however, is not governed solely by DNA repair. Local radiotherapy can initiate type I interferon-dependent innate and adaptive antitumor immunity, and STING-dependent cytosolic DNA sensing is required for this effect in immunogenic tumor models. Subsequent studies showed that mitotic progression after DNA damage generates micronuclei that recruit cGAS, thereby converting genome damage into inflammatory signaling, while the exonuclease TREX1 degrades radiation-induced cytosolic DNA and can suppress tumor immunogenicity in a dose-dependent manner. Together, these findings place the handling of radiation-damaged DNA at the center of the transition from genotoxic stress to immune activation ^[5–9]^. A related layer of therapeutic relevance is immunogenic cell death (ICD). Surface exposure of calreticulin determines the immunogenicity of dying cancer cells and is specifically required for γ-irradiation-induced apoptosis. In parallel, ATP released from dying tumor cells activates P2X7/NLRP3 signaling in dendritic cells and supports IL-1β-dependent adaptive immunity. Radiation has also been shown to induce immunogenic modulation, including enhanced antigen processing and calreticulin exposure, thereby increasing susceptibility of surviving tumor cells to T-cell-mediated killing^[10–13]^. Despite these advances, the tumor-intrinsic molecules that connect radioresistance, DNA damage adaptation, and the immunogenic consequences of irradiation remain incompletely defined. Unbiased proteomics offers a useful approach for identifying such regulators from clinical material. In chronic myeloid leukemia (CML), prior blood and bone marrow proteomic studies have already shown that proteome profiling can reveal clinically informative proteins associated with therapy resistance, disease stratification, and treatment response, supporting its value as a biomarker-discovery platform ^[14–16]^. One candidate of particular interest is XRRA1 (X-ray radiation resistance associated 1). XRRA1 was originally identified during an investigation of tumor-cell radioresponse in the colorectal cancer clone HCT116Clone2_XRR, in which it was reported to be down-regulated in an X-ray-resistant clone. The original cloning study further showed that XRRA1 expression changed after X-irradiation in HCT116 clones with distinct X-ray response phenotypes and proposed that XRRA1 might participate in the response of tumor and normal cells to X-radiation^[17]^. Despite this suggestive origin, whether XRRA1 directly contributes to radioresistance, DNA damage signaling, or radiation-induced immunogenic cell death has remained unclear. On this basis, the present study was designed to define the role of XRRA1 in the radiation response through integrated proteomic discovery, clinical validation, and mechanistic functional analysis.

## Methodology

### Isolation and estimation of CML proteins by using sodium dodecyl-sulfate polyacrylamide gel electrophoresis and bicinchoninic acid assay

Protein was isolated from peripheral blood samples of healthy controls and patients with chronic myeloid leukemia (CML) using RIPA buffer containing protease inhibitors. Protein concentration was measured by bicinchoninic acid (BCA) assay at 562 nm, and protein quality was assessed by SDS–PAGE ^[18]^.

### Protein profiling was analyzed using liquid chromatography-tandem mass spectrometry

Proteins isolated from control and CML peripheral blood samples were subjected to LC–MS/MS-based proteomic profiling. Ten representative protein isolates from each group were normalized, pooled in equal volume, and processed for mass spectrometric analysis ^[18,19]^.

### In-solution protein digestion before liquid chromatography-tandem mass spectrometry

Pooled protein samples were diluted in 50 mM NH₄HCO₃, reduced with dithiothreitol, alkylated with iodoacetamide, and digested overnight with trypsin at 37 °C. Digested peptides were acidified with 0.1% formic acid, vacuum-dried, centrifuged, and the supernatant was collected for LC–MS/MS analysis.

### Protein identification by liquid chromatography-tandem mass spectrometry

Peptides were separated on a BHE C18 UPLC column and analyzed on a Waters Synapt G2 Q-TOF instrument. Raw data were processed using MassLynx 4.1 and PLGS software with a decoy database strategy at a 1% false discovery rate; carbamidomethylation of cysteine was used as a fixed modification and methionine oxidation as a variable modification ^[18]^.

### Protein-protein interaction analysis of XRRA1 using STRING

The XRRA1-centered protein–protein interaction network was generated using STRING v12.0 with *Homo sapiens* selected as the reference organism. Known and predicted associations were used to identify functional interactors and enriched pathways ^[20]^.

### Co-expression analysis using GEPIA3 and enrichment analysis using Enrichr

Gene co-expression analysis was performed using GEPIA3 based on TCGA and GTEx datasets, and the resulting co-expressed genes were functionally annotated using Enrichr for Gene Ontology and pathway enrichment analyses ^[21,22]^.

### Immunohistochemistry (IHC)

Formalin-fixed, paraffin-embedded tissue sections were deparaffinized, rehydrated, subjected to antigen retrieval in Tris–EDTA buffer (pH 9), and incubated with primary and secondary antibodies. Signal was developed with DAB, counterstained with hematoxylin, dehydrated, cleared, and mounted for microscopic evaluation ^[23]^.

### Digital pathology-based interpretation of IHC staining

Digital interpretation of IHC images was performed using DeepLIIF, which was used for stain separation, cell segmentation, positive-cell detection, and H-score-based quantification. Positive cells were categorized as weak, moderate, or strong, and H-scores were calculated as: H-score = (1 × % weak) + (2 × % moderate) + (3 × % strong) ^[24]^.

### Cell culture and irradiation

Human U251, T47D, PR-PC3, and CO-SW620 cells were cultured in RPMI-1640 complete medium, whereas HEK293 and NIH3T3 cells were maintained in F199 complete medium. All cells were grown at 37 °C in a humidified incubator with 5% CO₂.

### Ionizing radiation

Cells were exposed to 4, 6, or 8 Gy ionizing radiation using an X-RAD 320 irradiator and harvested at the indicated time points for western blotting, immunoprecipitation, or immunostaining.

### Quantitative real-time PCR

Total RNA was extracted using TRIzol reagent and quantified by NanoDrop spectrophotometry. cDNA was synthesized using reverse transcription kits, and quantitative real-time PCR was performed using TaqMan chemistry on a QuantStudio 6 Flex system; β-actin was used as internal controls for mRNA and analyses, respectively and all probes mentioned in Table S1.

### Cell transfection

Cells were transfected with XRRA1-targeting siRNA (Dharmacon) or the corresponding non-targeting control siRNA using Lipofectamine RNAiMAX (Invitrogen) according to the manufacturer’s instructions and were used for downstream assays at the indicated time points.

### Cell viability assay

Cells were seeded in 96-well plates at 2.0 × 10³ cells per well, and viability was measured using the CellTiter-Glo luminescent assay according to the manufacturer’s protocol.

### Clonogenic assay

For colony formation assays, transfected cells were replated in 6-well plates at 1.0 × 10³ to 1.5 × 10³ cells per well, irradiated as indicated, and cultured for 10–15 days. Colonies were fixed, stained with 0.5% crystal violet, and imaged.

### Spheroid formation assay

Transfected cells were seeded in 24-well ultra-low attachment plates at 5.0 × 10³ cells per well and cultured under spheroid-forming conditions for 7–14 days. Spheroids were imaged using an Olympus IX51 microscope.

### Apoptosis analysis

Cells were seeded in 6-well plates, treated as indicated, and stained using an Annexin V-FITC apoptosis detection kit (Abcam). Both floating and adherent cells were collected, and apoptotic cells were quantified by flow cytometry.

### Western blot analysis

Whole-cell lysates were prepared in RIPA buffer and quantified by BCA assay. Equal amounts of protein were resolved on gradient polyacrylamide gels, transferred onto nitrocellulose membranes, incubated with primary and HRP-conjugated secondary antibodies, and detected by chemiluminescence. Band intensity was quantified using ImageJ. All the antibodies’ details are in Table S2.

### Cell-cycle analysis

Cells were seeded in 6-well plates, transfected and treated as indicated, fixed after 72 h, and stained with FxCycle PI/RNase staining solution. DNA content was analyzed using a BD FACSCalibur flow cytometer.

### Immunofluorescence

Cells grown on coverslips were fixed with 4% paraformaldehyde, permeabilized with 0.2% Triton X-100, blocked in 3% BSA, and incubated with primary and Alexa Fluor-conjugated secondary antibodies. Nuclei were counterstained with DAPI, and images were acquired using a Zeiss LSM780 Elyra SIM microscope.

### Immunoprecipitation

Treated cells were lysed in TNN buffer containing protease inhibitors, and equal amounts of protein were pre-cleared with protein A/G agarose and IgG. Samples were incubated with the indicated antibodies overnight, immune complexes were captured with protein A/G agarose, and the eluted proteins were analyzed by western blotting with whole-cell lysates used as input controls.

### Immunogenic cell death assay

To evaluate immunogenic cell death, U251, T47D, PR-PC3, and CO-SW620 cells were seeded in 6-well plates and treated under the indicated conditions (control, si-XRRA1, 4 Gy, and si-XRRA1 + 4 Gy). Extracellular ATP release was measured in culture supernatants, and cell-surface calreticulin exposure was analyzed by flow cytometry using Alexa Fluor 488-conjugated anti-calreticulin antibody with the corresponding isotype control.

### Statistical analysis

Data are presented as mean ± SD from at least three independent experiments. Statistical comparisons were performed using an unpaired two-tailed Student’s t-test or one-way ANOVA with appropriate post hoc testing, and P < 0.05 was considered statistically significant.

### Bioinformatics tools

TIMER2.0 was used for tumor immune infiltration and gene-expression association analysis ^[25]^, GTEx for normal tissue expression profiling ^[26]^, ARCHS4 for transcriptomic/co-expression exploration^[27,28]^, and DepMap for gene dependency and codependency analyses across cancer cell lines ^[29]^.

## Results

### Integrated proteomic, transcriptomic and dependency analyses prioritize XRRA1 as a clinically relevant candidate in chronic myeloid leukemia

To identify candidate molecules associated with chronic myeloid leukemia (CML), we profiled peripheral blood from healthy controls and patients with CML by LC–MS/MS and then integrated these findings with transcriptomic, network, prognostic, and functional-dependency datasets as explained in Figure 1A–C; Supplementary Figure 1A–I; and in Tables S3–S8. Proteomic profiling detected 428 proteins in healthy controls and 345 proteins in CML, with 257 proteins shared between the two groups, 171 proteins unique to controls, and 88 proteins unique to CML (Fig. 1A). XRRA1 localized to the shared fraction, indicating that it was not completely lost in CML, but its proteomic representation was consistently reduced in the leukemia samples. Specifically, in healthy controls XRRA1 was detected with a PLGS score of 156.72, 30 peptides, 44.57% sequence coverage, 189 matched product ions and 2 modified peptides, whereas in CML the corresponding values declined to 90.58, 17 peptides, 16.79% coverage, 106 matched product ions and 1 modified peptide, while the theoretical peptide number remained unchanged at 69 in both datasets (Tables S3 and S4). Thus, relative to controls, XRRA1 showed an approximately 42% decrease in PLGS score, 43% fewer identified peptides, 62% lower sequence coverage and 44% fewer product ions in CML, nominating XRRA1 as a shared but quantitatively attenuated candidate in the disease proteome. To further determine whether this observation reflected a broader biological context, we next examined XRRA1 expression across tumor, normal tissue, tissue-atlas and cell-line datasets (Supplementary Fig. 1A–H). Pan-cancer comparison revealed that XRRA1 was not uniformly deregulated but showed recurrent, cancer-type-specific changes. Compared with matched normal tissues, XRRA1 expression was significantly higher in BLCA, BRCA, CHOL, COAD, ESCA, HNSC, LIHC, LUAD, LUSC and STAD, while UCEC showed the opposite pattern, with lower tumor expression than normal tissue; differential expression was also evident between primary and metastatic melanoma (Supplementary Fig. 1A). In normal tissues, XRRA1 exhibited broad constitutive expression, with testis representing a clear high-expression outlier and lower-to-moderate expression across multiple reproductives, epithelial, gastrointestinal, neural and stromal tissues (Supplementary Fig. 1B). Single-cell and tissue-resolved maps further indicated that XRRA1 is distributed across diverse cellular compartments rather than being restricted to one lineage, including epithelial, fibroblast/mesenchymal, muscle, neuronal and immune-associated populations (Supplementary Fig. 1C). This breadth was reinforced by system-level expression maps, which placed XRRA1 across connective, digestive, immune, integumentary, muscular, nervous, respiratory, urogenital/reproductive and cardiovascular systems, notably including bone marrow, lymphoid, myeloid, spleen and thymic compartments relevant to hematological disease (Supplementary Fig. 1G). A similar pattern was observed across cultured cell models, where XRRA1 expression was retained in a wide range of cell-line lineages, including lung (for example A549 and H1299), lymphoid/hematopoietic (Jurkat, HL60 and RAJI), breast/mammary (MCF7, MCF10A and MDAMB231), prostate (LNCaP, PC3 and DU145), kidney (293T and HEK293) and ovarian (SKOV3 and OVCAR3) backgrounds (Supplementary Fig. 1H). Together, these data indicate that XRRA1 is broadly expressed across normal tissues and transformed models but is differentially modulated across malignant contexts. Consistent with this broad expression pattern, transcript-structure analyses suggested that XRRA1 expression is supported by a largely conserved splice architecture rather than by highly tissue-restricted exon or isoform switching. Exon-level heatmaps showed coordinated read distribution across most annotated exons of the XRRA1 locus across tissues (Supplementary Fig. 1D), and junction-level maps similarly demonstrated recurrent, conserved splice connectivity across multiple tissue types (Supplementary Fig. 1E). Isoform analysis identified a dominant transcript, ENST00000321448.12, present across essentially all tissues, together with a smaller number of lower-abundance isoforms, including ENST00000531849.5 (Supplementary Fig. 1F). These observations argue that tissue-to-tissue variation in XRRA1 abundance is driven principally by changes in overall transcript level rather than wholesale changes in isoform identity. To place XRRA1 in functional context, we next analyzed XRRA1-associated pathways and interaction networks. Reactome enrichment of XRRA1 co-expressed genes highlighted pathways linked to post-transcriptional silencing by small RNAs (P = 5.11 × 10⁻⁴), regulation of NPAS4 mRNA translation (P = 1.09 × 10⁻³), regulation of CDH11 mRNA translation by microRNAs (P = 1.32 × 10⁻³), reversible hydration of carbon dioxide (P = 1.58 × 10⁻³), regulation of NPAS4 gene expression (P = 2.17 × 10⁻³), competing endogenous RNAs regulating PTEN translation (P = 4.00 × 10⁻³), regulation of MITF-M-dependent genes involved in apoptosis (P = 4.00 × 10⁻³), regulation of RUNX1 expression and activity (P = 8.01 × 10⁻³), TGFBR3 expression (P = 8.6 × 10⁻³) and regulation of PTEN mRNA translation (P = 8.6 × 10⁻³) (Fig. 1A). STRING network analysis further positioned XRRA1 as the hub of an 11-edge interaction network (Fig. 1B; Table S5). XRRA1 showed direct interactions with XRCC5, ADGRL1, ADGRL2, ADGRL3, GPALPP1, SPCS2, CLLU1OS, CHRDL2, SLFN5 and MAN2C1, while SLFN5 and CLLU1OS formed an additional secondary interaction. The strongest interaction was XRRA1-XRCC5 (score = 0.774), followed by XRRA1-ADGRL1, XRRA1-ADGRL2 and XRRA1-ADGRL3 (0.514 each), XRRA1-GPALPP1 (0.485), XRRA1-SPCS2 (0.474), XRRA1-CLLU1OS (0.447), XRRA1-CHRDL2 (0.418), XRRA1-SLFN5 (0.409) and XRRA1-MAN2C1 (0.405). Notably, the prominence of XRCC5, a core component of non-homologous end joining, is consistent with the annotated role of XRRA1 in radiation-response biology.

**Figure 1:**
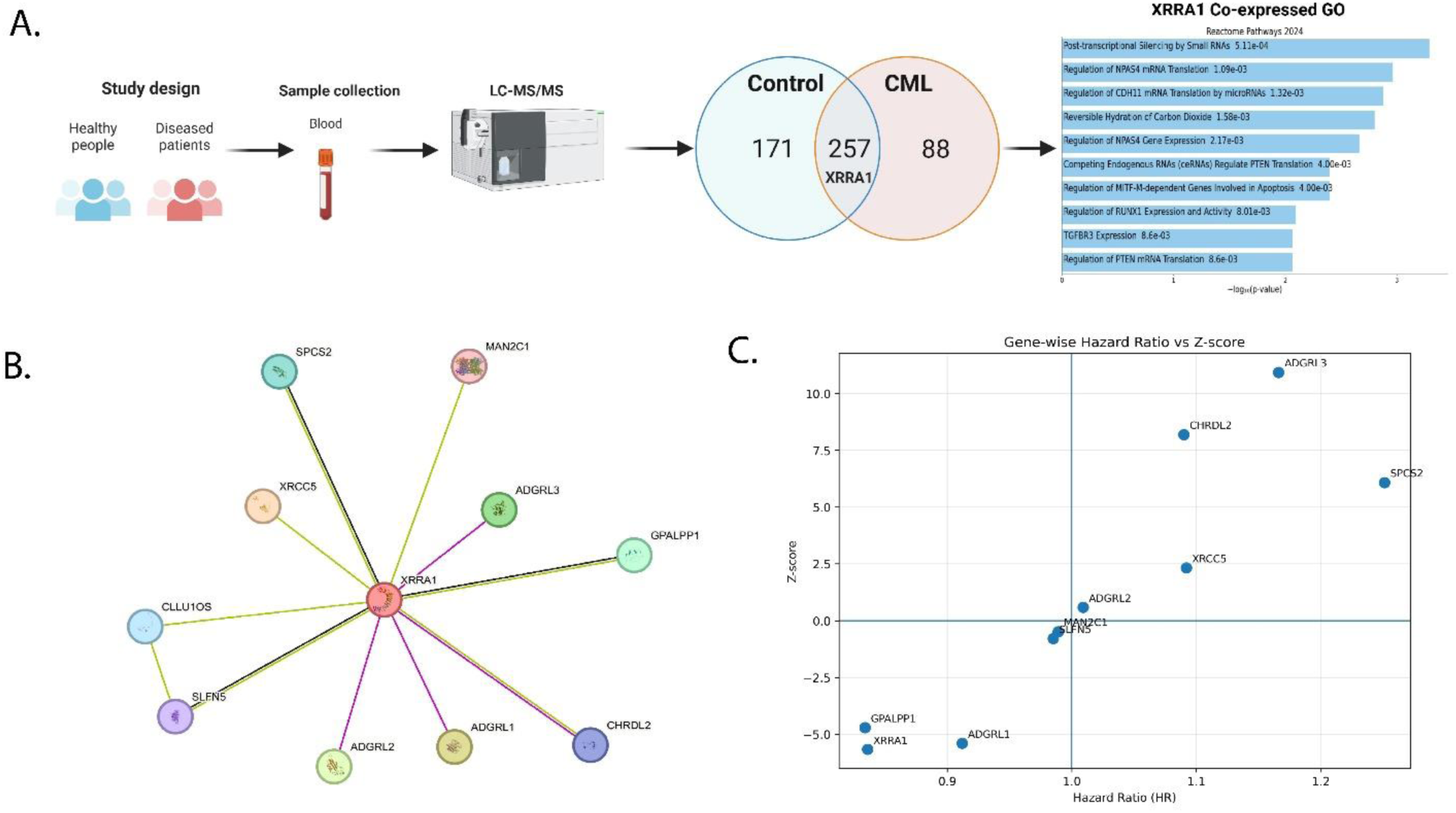
Proteomic discovery and systems-level prioritization of XRRA1 in chronic myeloid leukemia. **A,** Schematic overview of the study design and analytical workflow. Peripheral-blood samples from healthy controls and patients with chronic myeloid leukemia (CML) were subjected to LC–MS/MS profiling, followed by non-redundant protein overlap analysis and pathway interrogation. The Venn diagram shows proteins uniquely detected in controls (171), shared between control and CML samples (257), and uniquely detected in CML (88), with XRRA1 present in the shared fraction. The bar plot shows the top Reactome pathways enriched among XRRA1-associated genes, including post-transcriptional silencing by small RNAs, regulation of NPAS4 mRNA translation, regulation of CDH11 mRNA translation by microRNAs, reversible hydration of carbon dioxide, regulation of NPAS4 gene expression, ceRNA-mediated regulation of PTEN translation, regulation of MITF-M-dependent genes involved in apoptosis, regulation of RUNX1 expression and activity, TGFBR3 expression, and regulation of PTEN mRNA translation. Bar length indicates −log10(*P*). **B,** STRING protein–protein interaction network centered on XRRA1. XRRA1 is connected to XRCC5, ADGRL1, ADGRL2, ADGRL3, GPALPP1, SPCS2, CLLU1OS, CHRDL2, SLFN5 and MAN2C1, with an additional SLFN5-CLLU1OS interaction. **C,** Gene-wise hazard ratio (HR) versus Z-score plot for XRRA1 and XRRA1-associated genes. The vertical line indicates HR = 1, and the horizontal line indicates Z = 0. Genes in the lower-left quadrant are associated with favorable outcome, whereas genes in the upper-right quadrant are associated with increased risk. Numerical survival statistics are provided in Supplementary Table 6. Data underlying a are provided in Supplementary Tables 3 and 4, and data underlying b are provided in Supplementary Table 5.

We then asked whether the XRRA1-centred network carried clinical significance. In gene-wise hazard ratio analysis, XRRA1 localized to the favorable prognostic quadrant and was associated with reduced risk (HR = 0.836, 95% CI 0.785–0.889, Z = -5.665, P = 1.47 × 10⁻⁸) (Fig. 1C; Table S6). Similar protective associations were observed for GPALPP1 (HR = 0.834, P = 2.47 × 10⁻⁶) and ADGRL1 (HR = 0.912, P = 6.65 × 10⁻⁸). By contrast, ADGRL3 (HR = 1.166, P = 8.43 × 10⁻²⁸), CHRDL2 (HR = 1.090, P = 2.59 × 10⁻¹⁶), SPCS2 (HR = 1.251, P = 1.21 × 10⁻⁹) and XRCC5 (HR = 1.092, P = 0.0202) were associated with poorer outcome, whereas ADGRL2, MAN2C1 and SLFN5 were not significantly associated with survival (Table S6). Finally, DepMap codependency analysis showed that XRRA1 also resides within a distinct functional dependency landscape (Supplementary Fig. 1I; Tables S7 and S8). In the CRISPR dataset, the strongest positive associations were OR4D9 (r = 0.215), ADAM11 (r = 0.213), CAMKK2 (r = 0.213), MOV10 (r = 0.198) and DRD4 (r = 0.190), whereas the strongest negative associations were VWA5A (r = -0.196), HLA-DQB1 (r = -0.192), HLA-G (r = -0.190), KRTAP4-6 (r = -0.190) and ACSM2B (r = -0.190) (Table S7). In the RNAi dataset, the strongest positive associations were LGALS16 (r = 0.428), DBR1 (r = 0.427), CFAP52 (r = 0.421), RBX1 (r = 0.418) and CT45A1 (r = 0.415), whereas CYTH3 (r = -0.461), GAL (r = -0.397), RP1L1 (r = -0.391), RPS24 (r = -0.385) and LINC01559 (r = -0.374) were the strongest negative associations (Table S8). As shown in Supplementary Fig. 1I, the positively associated CRISPR and RNAi gene sets, and likewise the negatively associated gene sets, showed no cross-platform overlap, emphasizing strong platform-specific context dependence; notably, GSTK1 was the only gene shared between Tables S7 and S8, and it switched directionality between platforms. Pathway enrichment of the positively associated genes pointed to S/G2 phase transition, G2/M phase transition, male sex determination, AMPK signaling, EGFR/ERBB3→MEF/MYOD/NFATC/MYOG signaling and the SCF/CCNF complex, whereas negatively associated genes enriched direct DNA repair, direct DNA repair suppression in cancer, ganglioside-type glycosphingolipid biosynthesis and cilia disorganization (Supplementary Fig. 1I). Collectively, these data identify XRRA1 as a broadly expressed, structurally conserved and clinically favorable candidate whose diminished proteomic representation in CML occurs within a network linked to RNA regulation, cell-cycle control and DNA-damage-associated biology.

### XRRA1 protein expression is enriched in radiation-exposed clinical tissues

To validate XRRA1 at the protein level in human specimens, we examined a clinicopathologically annotated cohort of 21 biopsy samples by conventional immunohistochemistry, DeepLIIF-assisted digital pathology, and immunoblotting (Fig. 2; Supplementary Fig. 2; Tables S9 and S10). Cohort-wide ranking of XRRA1 staining revealed a clear segregation by radiation exposure history. Notably, all 18 CT/X-ray–exposed specimens occupied the top 18 positions in the XRRA1 H-score distribution, with values ranging from 100.3 to 194.3 and a median of 142.6, whereas the three non-exposed specimens formed the bottom of the distribution, with H-scores of 91.7, 88.6, and 6.2 (median 88.6) (Table S9). This separation was also reflected in staining intensity patterns. The top-ranked case (LL320L-24) showed 84.1% IHC-positive cells with 220 weak (1+), 449 moderate (2+), and 620 strong (3+) cells, whereas the lowest-ranked case (LL184L-24) showed only 4.9% positivity with 68 weak, 17 moderate, and 3 strong cells, indicating a broad dynamic range of XRRA1 expression across the clinical cohort. Representative images confirmed that XRRA1 staining was stronger in radiation-exposed tissue than in tissue with no radiation history, approaching the intense signal observed in placenta, which served as a positive control (Fig. 2A). This pattern was preserved across the original bright-field IHC image and the corresponding XRRA1, DAPI, hematoxylin, Lamin-associated polypeptide-2 (LAP2), and segmentation panels, supporting the specificity of the differential signal. DeepLIIF-based quantification of the representative fields further substantiated this finding (Table S10). The radiation-exposed specimen contained 1,458 XRRA1-positive cells among 1,895 nuclei analyzed (76.9%), with an H-score of 122.8, whereas the no-radiation specimen contained 1,086 positive cells among 1,845 nuclei (58.9%), with an H-score of 73.8. Importantly, the exposed tissue also showed a clear enrichment of higher-intensity staining classes, containing 458 moderate and 206 strong cells compared with 226 moderate and only 25 strong cells in the non-exposed tissue. As expected, the placenta positive control showed robust XRRA1 positivity (1,101/1,473 nuclei; 74.7%; H-score 128.0), whereas the negative control remained minimal (85/2,546 nuclei; 3.3%; H-score 3.5). Supplementary Fig. 2 extended these observations across representative exposure categories and suggested that XRRA1 abundance tracks with prior radiation burden. A specimen from a patient after a second radiation therapy cycle showed the strongest signal, with 1,719 nuclei analyzed, 67.9% XRRA1-positive cells, and an estimated H-score of 225.9. This exceeded the representative specimen obtained after a first radiation therapy cycle (1,765 nuclei; 52.5% positive; H-score 93.3) and the specimen from a patient with no radiation therapy (2,430 nuclei; 47.5% positive; H-score 62.7). The corresponding positive and negative controls yielded estimated H-scores of 180.1 and 0.0, respectively, confirming the dynamic range of the assay. Further RT-PCR results showed that XRRA1/ACTN ratios were significantly higher in radiation-exposed tumor samples than in either healthy controls or tumor samples without radiation history (Fig. 2B). Orthogonal protein validation by immunoblotting confirmed XRRA1 at the expected ∼90 kDa molecular weight relative to ACTB (42 kDa) in both healthy and patient-derived samples (Fig. 2C). Together, these data establish XRRA1 as a radiation-associated protein that is consistently elevated in clinically irradiated human tissues, as demonstrated by cohort-level IHC ranking, digital H-score analysis, and immunoblot validation.

**Figure 2:**
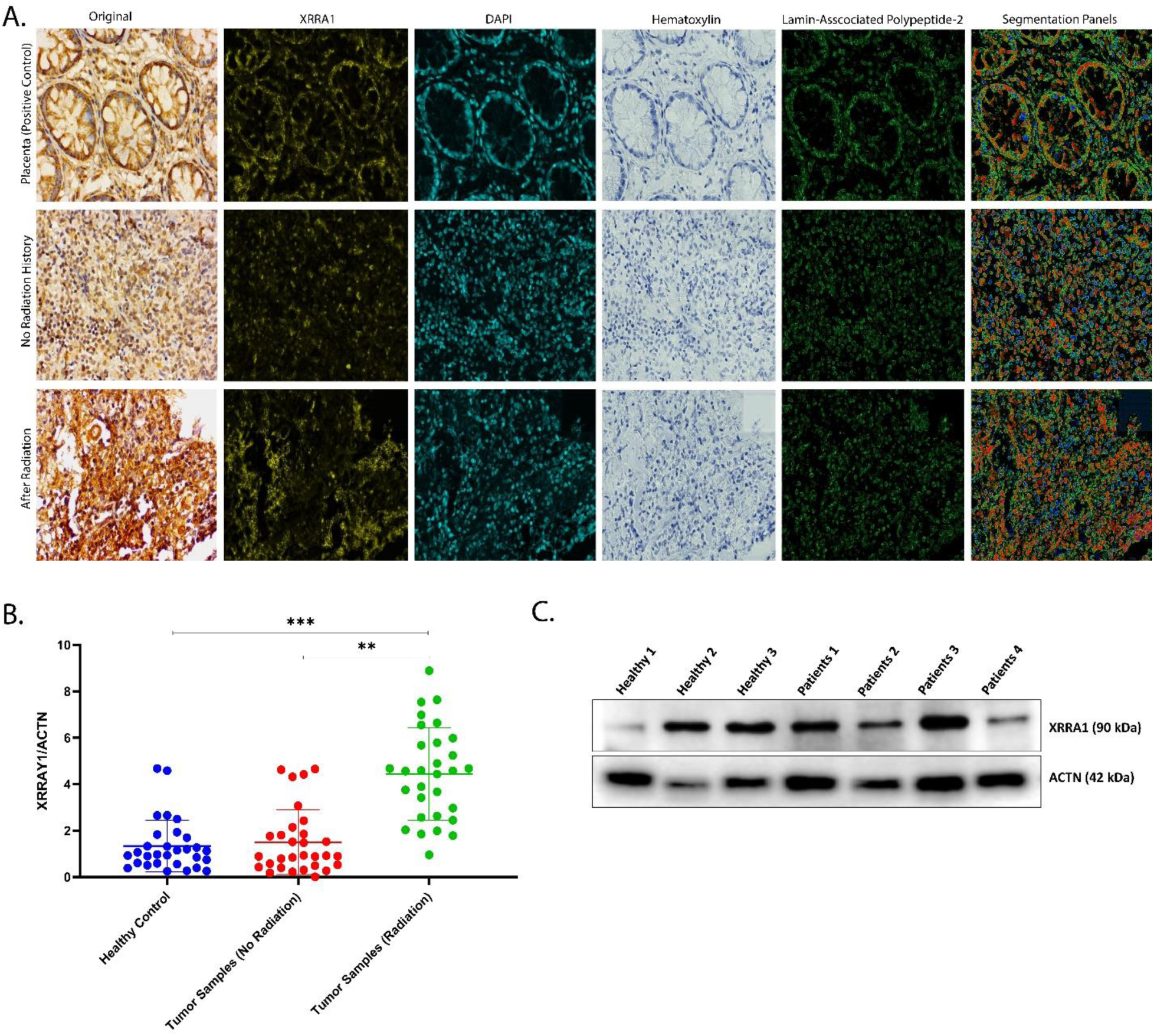
Clinical validation of XRRA1 protein expression in radiation-exposed human tissues. **A,** Representative XRRA1 immunohistochemistry and DeepLIIF-based image decomposition in placenta (positive control), tumor tissue without radiation history, and radiation-exposed tumor tissue. For each specimen, the original bright-field image and the corresponding XRRA1, DAPI, hematoxylin, LAP2, and segmentation panels are shown. **B,** Densitometric quantification of XRRA1/ACTN immunoblot signal in healthy controls, tumor samples without radiation history, and radiation-exposed tumor samples. Each dot represents an individual sample. Statistical comparisons are indicated in the plot. **C,** Representative immunoblot showing XRRA1 (∼90 kDa) and ACTN (∼42 kDa) in healthy and patient-derived samples used for protein-level validation.

### XRRA1 is robustly induced by ionizing radiation in both normal & cancer cells

To determine whether XRRA1 is dynamically regulated by ionizing radiation, normal and cancer cells were exposed to a single 4 Gy dose and harvested over a 2-72 h time course. Immunoblot analysis showed that XRRA1 was radiation responsive in both cellular contexts, but the magnitude of induction differed markedly between them (Fig. 3A). In normal cells, XRRA1 increased rapidly after irradiation and remained elevated throughout the time course, with particularly strong accumulation at later time points. By contrast, cancer cells showed only a comparatively moderate increase in XRRA1 after irradiation in compared to control. PARP expressions were also altered over time in both groups, consistent with activation of a radiation-associated stress response, whereas ACTN served as the loading control. Further RT-PCR analysis confirmed that the XRRA1 response to radiation was substantially greater in normal cells than in cancer cells (Fig. 3B). Relative to baseline, XRRA1 in normal cells rose to approximately 2.8-fold at 2 h, remained elevated at intermediate time points, and increased further to roughly 4.3-fold at 48 h and 6.4-fold at 72 h. In contrast, cancer cells displayed a blunted response, with XRRA1 remaining within an approximately 1.5–2.5-fold range across the same interval and reaching only about 2.5-fold at 48–72 h. Thus, although XRRA1 is inducible by radiation in both cell types, the amplitude and persistence of induction are far more pronounced in normal cells. We next examined whether XRRA1 expression carried clinical relevance. Kaplan–Meier analysis stratified by XRRA1 expression (cutoff = 8.680) showed a highly significant separation in survival probability (P = 1.088 × 10⁻¹⁴) (Fig. 3C). In keeping with the earlier hazard-ratio analysis, the overall trend of the survival curves indicated that higher XRRA1 expression was associated with a more favorable outcome over most of the follow-up period. Finally, qualitative assessment of post-irradiation cell growth revealed distinct phenotypic responses in normal and cancer cells (Fig. 3D). In normal cells, irradiated cultures retained dense adherent growth and appeared more confluent than the non-irradiated condition. In contrast, cancer cells showed the opposite trend: non-irradiated cultures formed larger, compact cellular aggregates, whereas irradiated cultures exhibited reduced colony burden with fewer and smaller residual cell clusters. Together, these findings identify XRRA1 as a radiation-inducible protein with stronger induction in normal cells, while also linking higher XRRA1 expression to better patient survival and to differential post-irradiation growth behaviors in normal versus malignant cells.

**Figure 3:**
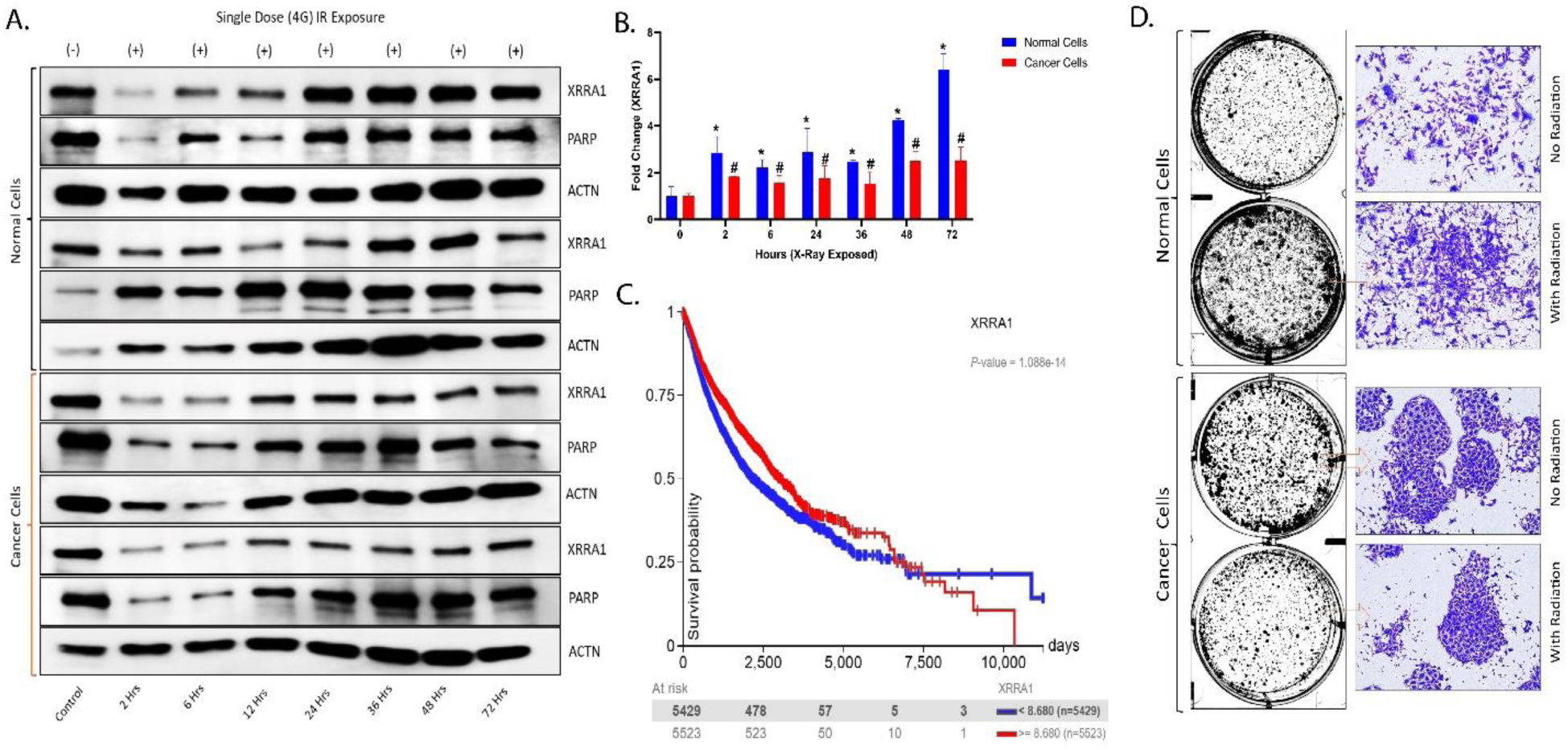
XRRA1 is induced by ionizing radiation with distinct kinetics in normal and cancer cells and is associated with survival outcome. **A,** Representative immunoblots showing XRRA1 and PARP expression in normal and cancer cells collected at the indicated time points after a single 4 Gy (G) ionizing radiation exposure (0, 2, 6, 12, 24, 36, 48 and 72 h). ACTN was used as the loading control. Two representative immunoblot series are shown for each cell type. **B,** Densitometric quantification of XRRA1 fold changes in normal cells (blue) and cancer cells (red) after irradiation, with expression at 0 h set to 1. Error bars indicate variability across measurements. Statistical significance is indicated in the plot. **C,** Kaplan–Meier survival analysis stratified by XRRA1 expression using a cutoff value of 8.680. Blue line, XRRA1 < 8.680; red line, XRRA1 ≥ 8.680. The *P* value is shown in the panel. **D,** Representative whole-well and higher-magnification images showing the growth pattern of normal and cancer cells under no-radiation and radiation conditions. Boxed regions indicate the areas shown at higher magnification.

### XRRA1 depletion potentiates radiation-induced apoptotic cell death across multiple cancer cell lines

To determine whether XRRA1 functionally contributes to cellular resistance to irradiation, we silenced XRRA1 using an ON-TARGET plus SMART pool comprising four independent human XRRA1-targeting siRNAs (Supplementary Fig. 3A; Supplementary Table 11). The selected reagent was predicted to target 69/69 annotated XRRA1 accessions, supporting broad transcript coverage. XRRA1 silencing was efficient in all tested models, reducing XRRA1 expression to approximately 6–10% of control levels in U251, BR-T47D, PR-PC3 and CO-SW620 cells (Supplementary Fig. 3B), thereby establishing a robust system for functional interrogation. We next assessed the effect of XRRA1 loss on cell survival following irradiation. Across all four cancer cell lines, XRRA1 knockdown alone had only a minimal effect on basal viability, with values remaining close to control. Exposure to 4 Gy alone produced a moderate reduction in viability, lowering survival to roughly 73–86% depending on the cell line. By contrast, the combination of si-XRRA1 and 4 Gy caused a marked loss of viability in every model tested, reducing cell survival to approximately 31% in U251, 35% in BR-T47D, 32% in PR-PC3, and 26% in CO-SW620 cells (Fig. 4A). Thus, although XRRA1 depletion by itself was not strongly cytotoxic, it substantially enhanced the anti-proliferative effect of radiation. To determine whether this loss of viability reflected apoptotic cell death, we quantified caspase 3/7 activity under the same treatment conditions. In all four cell lines, control and si-XRRA1-alone groups remained near baseline, whereas 4 Gy alone induced a modest increase in caspase activity, ranging from approximately 1.5-fold to 2.6-fold. Notably, combined XRRA1 silencing plus irradiation triggered a much stronger caspase response, reaching roughly 5.9-fold in U251, 6.8-fold in BR-T47D, 7.9-fold in PR-PC3, and 5.8-fold in CO-SW620 cells (Fig. 4B). These data indicate that loss of XRRA1 markedly amplifies radiation-induced executioner caspase activation. Flow-cytometric Annexin V-FITC/PI analysis further confirmed that the combination treatment increased apoptotic cell death (Fig. 4C). In the representative analysis, the live-cell fraction (Q4) decreased from 93.7% in the control group to 73.1% after si-XRRA1 + 4 Gy, while the apoptotic fractions expanded substantially. Specifically, the late apoptotic population (Q2) increased from 2.33% in control cells to 17.0% after combination treatment, and the early apoptotic population (Q3) increased from 3.60% to 8.97%. When early and late apoptotic cells were combined, the total Annexin V-positive fraction increased from 5.93% in control cells and 6.55% after si-XRRA1 alone to 10.65% after 4 Gy alone and 25.97% after combined treatment. Quantification of apoptotic cells across replicate measurements showed the same trend, with apoptosis rising from approximately 6-7% in control and si-XRRA1 groups to ∼13-14% after radiation alone and ∼28% after combined XRRA1 silencing and irradiation (Supplementary Fig. 3C). Together, these findings identify apoptosis as a major mechanism underlying the radiosensitizing effect of XRRA1 loss. To place these functional observations in a signaling context, kinase enrichment analysis (KEA) was performed for XRRA1-associated interactions (Supplementary Table 12). This analysis ranked PDK4 and PDK3 as the top predicted kinase associations with XRRA1 (Z-score = 8.22 each), followed by TESK1 (6.29), FRK (5.74), PDK2 (5.31) and PRKD3 (4.97). These predictions support the view that XRRA1 may participate in stress-response and survival-associated kinase networks that become functionally important during radiation exposure. Collectively, the data indicates that XRRA1 acts as a protective determinant against radiation-induced apoptosis, and that its depletion markedly sensitizes tumor cells to irradiation.

**Figure 4:**
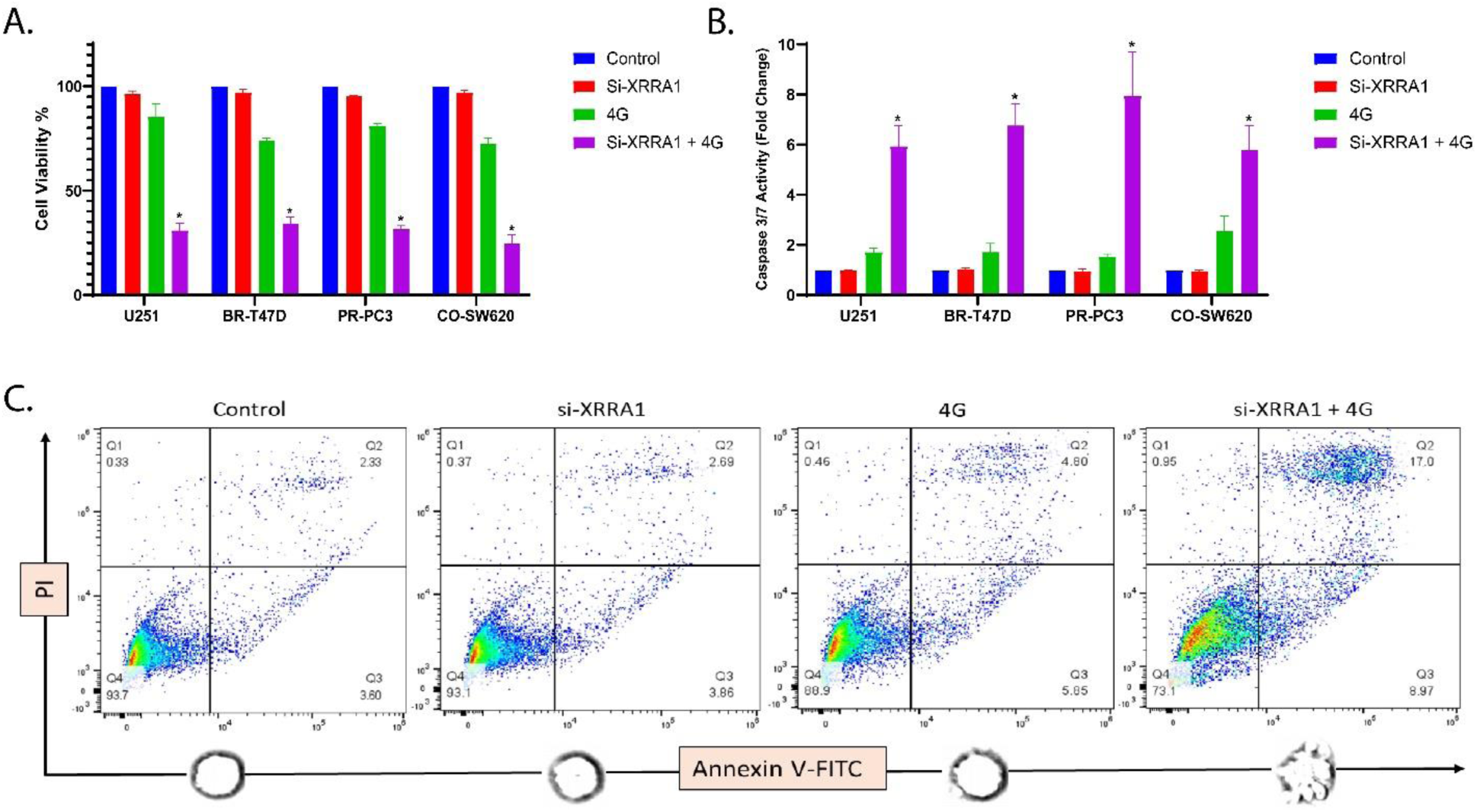
XRRA1 depletion sensitizes cancer cells to radiation-induced loss of viability and apoptosis. **A,** Relative cell viability in U251, BR-T47D, PR-PC3 and CO-SW620 cells under control, si-XRRA1, 4 Gy, and si-XRRA1 + 4 Gy conditions. XRRA1 silencing alone had limited effect on basal viability, whereas combined XRRA1 depletion and irradiation markedly reduced viability in all four cell lines. **B,** Caspase-3/7 activity measured under the same treatment conditions shown in a. Combined si-XRRA1 + 4 Gy induced a substantially greater increase in caspase activity than either treatment alone across all cell lines. **C,** Representative Annexin V–FITC/PI flow-cytometry plots for the indicated treatment groups. Quadrants denote Q1, PI-positive/Annexin V-negative; Q2, Annexin V-positive/PI-positive; Q3, Annexin V-positive/PI-negative; and Q4, double-negative viable cells. Numbers indicate the percentage of cells in each quadrant. The combination of XRRA1 silencing and irradiation increased both early and late apoptotic populations. Related knockdown validation and apoptosis quantification are shown in Supplementary Fig. 3, and siRNA details are provided in Supplementary Table 11.

### XRRA1 depletion suppresses clonogenic recovery and intensifies radiation-associated DNA damage signaling

To determine whether the loss of short-term viability observed after XRRA1 silencing translated into impaired long-term survival, we examined spheroid growth, colony formation, and DNA damage signaling following XRRA1 knockdown and irradiation (Fig. 5; Supplementary Fig. 4; Supplementary Table 13). In spheroid assays, control cells formed large, compact aggregates, whereas either si-XRRA1 or 4 Gy alone caused only partial disruption. By contrast, the combined si-XRRA1 + 4 Gy condition markedly reduced spheroid integrity, leaving only small residual aggregates (Fig. 5A). A similar pattern was observed in 2D colony-foci assays, in which combined XRRA1 depletion and irradiation produced a pronounced collapse in colony density relative to either single treatment (Fig. 5B). These findings indicate that XRRA1 supports both anchorage-independent and anchorage-dependent recovery after radiation stress. This phenotype was reproduced across multiple tumor cell models in dose-escalation clonogenic assays. In U251, CO-SW620, T47D, and PR-PC3 cells, radiation alone reduced colony outgrowth, but this effect was consistently enhanced by XRRA1 silencing (Supplementary Fig. 4A–D). In all four models, the addition of 4 Gy to si-XRRA1 further diminished colony number and size, and escalation to 6 Gy and 8 Gy resulted in progressively greater loss of surviving colonies, with only sparse residual growth in the higher-dose conditions. Thus, XRRA1 appears to sustain clonogenic recovery over a broad range of tumor contexts.

**Figure 5:**
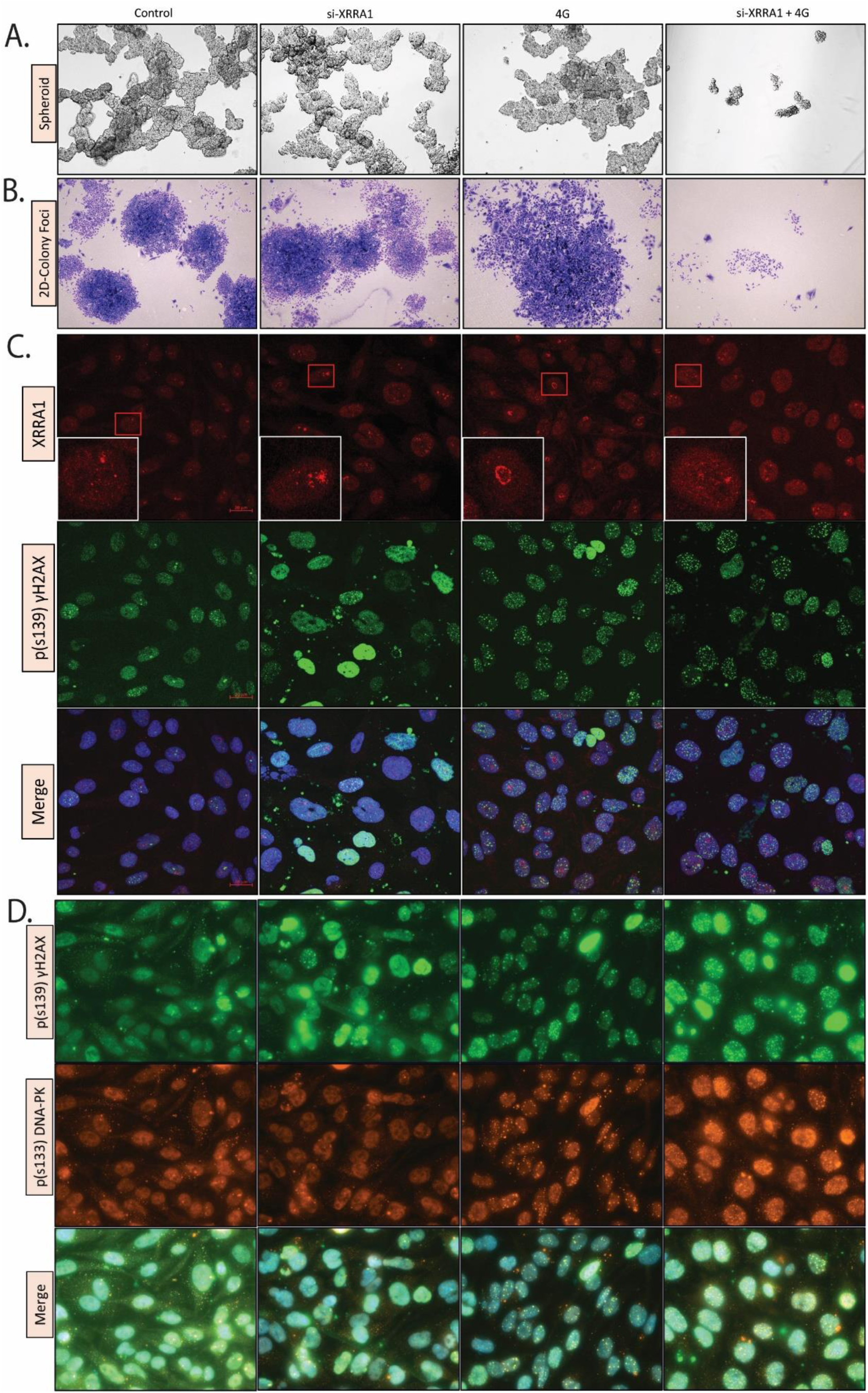
XRRA1 depletion disrupts spheroid and colony growth and enhances radiation-associated DNA damage signaling. **A,** Representative bright-field images of spheroid growth under control, si-XRRA1, 4 Gy, and si-XRRA1 + 4 Gy conditions. Combined XRRA1 silencing and irradiation caused the greatest loss of spheroid integrity. **B,** Representative images of 2D colony-foci formation under the same treatment conditions shown in a. Colony-foci density was most strongly reduced in the si-XRRA1 + 4 Gy group. **C,** Immunofluorescence images showing XRRA1 (red), p(S139)γH2AX (green), and merged images with nuclear counterstain (blue) under the indicated treatment conditions. Boxed regions are shown at higher magnification in the inset panels. **D,** Immunofluorescence images showing p(S139)γH2AX (green), p(S133) DNA-PK (orange), and merged images under the indicated treatment conditions. Combined XRRA1 depletion and irradiation produced prominent DNA damage-associated nuclear puncta. Related dose-escalation and quantitative microscopy data are shown in Supplementary Fig. 4 and Supplementary Table 13.

We next asked whether this reduction in survival was accompanied by enhanced DNA damage signaling. Immunofluorescence analysis showed that p(S139) γH2AX nuclear puncta increased after either XRRA1 depletion or irradiation and became more extensive when both perturbations were combined (Fig. 5C). In parallel, staining for p(S139) γH2AX and p(S133) DNA-PK revealed stronger dual nuclear puncta after irradiation, with the most prominent signal observed when XRRA1 knockdown was paired with radiation (Fig. 5D). Supplementary dose-escalation imaging confirmed that γH2AX accumulation intensified further as radiation dose increased from 4 Gy to 6 Gy and 8 Gy in XRRA1-depleted cells (Supplementary Fig. 4E). Quantitative image analysis supported these observations (Supplementary Table 13). In control cells, the mean background-subtracted γH2AX signal was 17.20, with 5.16% γH2AX-positive nuclei. After 4 Gy, this increased to 49.34 and 61.63% positive nuclei (2.87-fold over control). Notably, si-XRRA1 alone increased γH2AX even more strongly, to 74.84 with 62.93% positive nuclei (4.35-fold over control), indicating that XRRA1 depletion itself imposes basal genomic stress. In XRRA1-depleted cells exposed to escalating radiation doses, the γH2AX signal remained high at 4 Gy (46.09, 67.23% positive nuclei), increased further at 6 Gy (70.06, 85.18% positive nuclei), and peaked at 8 Gy (91.24, 94.43% positive nuclei; 5.30-fold over control). By contrast, quantitative comparison of the XRRA1 red channel should be interpreted cautiously, as the composite microscopy panels were display-normalized PNG images rather than raw acquisition files, making γH2AX the more reliable quantitative readout in this dataset. Finally, co-immunoprecipitation analysis extended these findings to DNA repair-associated proteins. Immunoblots from Ku70 and XRCC4 immunoprecipitated, together with corresponding input lysates from U251 and T47D cells under control and irradiated conditions, linked XRRA1 to proteins involved in the non-homologous end-joining machinery (Supplementary Fig. 4F). Taken together, these data identify XRRA1 as a determinant of post-irradiation survival whose loss compromises clonogenic recovery and is associated with persistent DNA damage signaling.

### XRRA1 loss cooperates with irradiation to engage cGAS-STING signaling and hallmarks of immunogenic cell death

Given the increase in DNA damage after XRRA1 depletion, we next asked whether loss of XRRA1 also promotes cytosolic DNA sensing and downstream immunogenic stress signaling. Immunoblot analysis performed under Control, si-XRRA1, 4 Gy, and si-XRRA1 + 4 Gy conditions showed the expected induction of XRRA1 by irradiation and its suppression by siRNA and further revealed activation of the cGAS–STING pathway (Fig. 6A). Relative to the single-treatment groups, the combination condition showed stronger STING and pS172-TBK1 signals together with sustained pS396-IRF3, consistent with enhanced signaling through the cGAS-STING-TBK1-IRF3 axis. cGAS was increased in irradiated cells, whereas HMGA2 was reduced after XRRA1 depletion, particularly in the irradiated setting. The same blot also showed altered abundance of multiple stress- and cell-cycle-associated regulators, including acetyl-p53, pS20-p53, pS392-p53, p16-INK4A, p14-ARF, CDK6 and CDK4, indicating broad remodeling of damage-response signaling after combined XRRA1 loss and radiation exposure. Cell-cycle profiling suggested that XRRA1 silencing alone had little effect on basal phase distribution, whereas irradiation redistributed cells out of S phase in both U251 and T47D cells (Fig. 6B). In U251 cells, the S-phase fraction decreased from approximately 27% in controls to about 12% after 4 Gy and remained low after si-XRRA1 + 4 Gy, with reciprocal enrichment of the G1 and G2/M compartments. In T47D cells, radiation similarly reduced the S-phase population and increased the G2/M fraction, and this redistribution was maintained under the combination condition. Thus, the dominant effect of XRRA1 depletion in irradiated cells was not the generation of a distinct new cell-cycle state, but rather an amplification of the radiation-driven stress response. This shift was accompanied by strong induction of interferon-stimulated genes. Quantitative expression analysis showed only modest changes in IFIT3, OAS2, BST2, IRGM, GBP2, STAT1, OAS1, IFI44 and ISG15 after si-XRRA1 alone, and a moderate induction after 4 Gy alone, generally in the ∼2–4-fold range (Fig. 6C). By contrast, combined XRRA1 depletion and irradiation produced a much stronger response, increasing these transcripts to approximately ∼5-9-fold above baseline, with the largest induction observed for IRGM, OAS2, OAS1 and IFI44. These findings place XRRA1 upstream of a radiation-triggered interferon-like transcriptional program. We next examined whether this signaling state was accompanied by features of immunogenic cell death. Extracellular ATP release remained near background in the Control and si-XRRA1 groups, increased modestly after 4 Gy, and rose strikingly after the combined treatment across all four cancer cell lines analyzed (U251, BR-T47D, PR-PC3 and CO-SW620) (Fig. 6D). ATP levels reached approximately 52–65 nM ATP ml⁻¹ per 10⁵ cells after si-XRRA1 + 4 Gy, compared with only ∼11–15 nM ATP ml⁻¹ per 10⁵ cells after 4 Gy alone. Consistently, flow-cytometric analysis of cell-surface calreticulin showed a progressive rightward shift from control to irradiation alone and a further increase after the combination treatment (Fig. 6E). The gated calreticulin-associated signal increased from 1.44 in control cells and 4.78 after si-XRRA1 alone to 15.2 after 4 Gy and 36.6 after si-XRRA1 + 4 Gy, whereas matched isotype controls remained comparatively low. Together, these data indicate that XRRA1 restrains radiation-induced cGAS–STING activation, interferon-stimulated gene expression, ATP release and surface calreticulin exposure. Loss of XRRA1 therefore converts the radiation response into a more immunogenic state.

**Figure 6:**
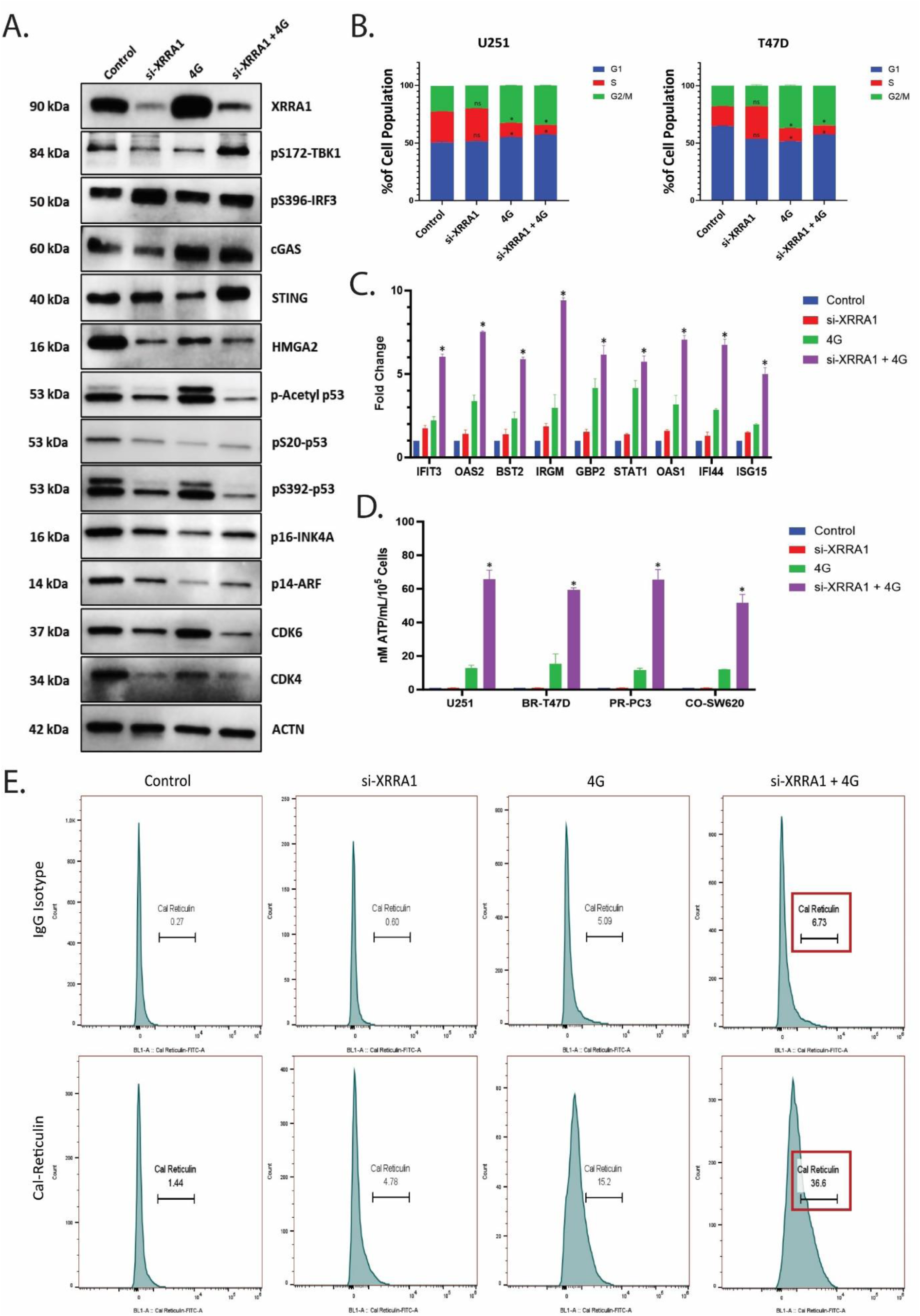
XRRA1 depletion enhances cGAS–STING signaling and immunogenic cell-death-associated outputs after irradiation. **A,** Representative immunoblots showing XRRA1, pS172-TBK1, pS396-IRF3, cGAS, STING, HMGA2, acetyl-p53, pS20-p53, pS392-p53, p16-INK4A, p14-ARF, CDK6, CDK4 and ACTN under control, si-XRRA1, 4 Gy, and si-XRRA1 + 4 Gy conditions. **B,** Stacked bar plots showing cell-cycle distribution (G1, S and G2/M) in U251 and T47D cells under the indicated treatment conditions. Radiation redistributed cells out of S phase, whereas si-XRRA1 alone had minimal effect. **C,** Fold-change expression of IFIT3, OAS2, BST2, IRGM, GBP2, STAT1, OAS1, IFI44 and ISG15 under control, si-XRRA1, 4 Gy, and si-XRRA1 + 4 Gy conditions. Combined XRRA1 depletion and irradiation produced the strongest induction of interferon-stimulated genes. **D,** Extracellular ATP release measured in U251, BR-T47D, PR-PC3 and CO-SW620 cells under the indicated treatment conditions. The y axis is expressed as nM ATP mL^−1^ per 10^*5^ cells. **E,** Representative flow-cytometry histograms showing cell-surface calreticulin staining under control, si-XRRA1, 4 Gy, and si-XRRA1 + 4 Gy conditions. The upper row shows matched IgG isotype controls, and the lower row shows anti-calreticulin staining. Values indicate the gated calreticulin-associated signal for each condition.

## Discussion

Radiotherapy is now understood as both genotoxic and immunologic therapy. In addition to generating lethal DNA lesions, ionizing radiation can trigger type I interferon-dependent tumor control through cytosolic DNA sensing and downstream innate immune activation. Against this background, the present study identifies XRRA1 as a previously underappreciated regulator of radioresponse, linking DNA damage adaptation, tumor survival, and radiation-induced immunogenicity. XRRA1 was originally described from a radiation-response model, but its significance in human cancer remained poorly resolved. By integrating CML discovery proteomics, systems-level prioritization, clinical tissue validation, and functional studies across multiple tumor models, our work positions XRRA1 at the center of a biologically and clinically meaningful radiation-response axis ^[5,6,17]^. The discovery arm of the study is important because it frames XRRA1 within a clinically relevant proteomic context. Prior blood proteomic studies in chronic myeloid leukemia have shown that proteomics can uncover biomarkers linked to disease state, treatment response, and resistance. In that setting, our identification of XRRA1 as a quantitatively diminished component of the CML proteome, yet one associated with a favorable prognostic signature, suggests that XRRA1 is not simply an oncogenic marker. Rather, our data support a context-dependent model in which baseline XRRA1 may reflect preserved cellular homeostasis or a less aggressive disease state, whereas under therapeutic genotoxic stress, XRRA1 behaves as an inducible survival factor that helps damaged cells adapt and recover ^[14–16]^. The clinical validation arm strengthens this interpretation. In the 21-biopsy cohort, all 18 CT/X-ray–exposed cases ranked above the three non-exposed cases by XRRA1 H-score, and digital pathology showed enrichment of moderate-to-strong XRRA1 staining in irradiated tissues. Together with immunoblot validation and representative positive-control placenta staining, these findings extend the radiation-associated identity of XRRA1 from experimental systems to human specimens and suggest that XRRA1 may function as a tissue-level marker of prior genotoxic exposure or adaptive radioresponse ^[17]^. The in vitro kinetic data further clarify this biology. Ionizing radiation induced XRRA1 in both normal and malignant cells, but the response was stronger and more sustained in normal cells. Functionally, XRRA1 knockdown alone had only a limited effect on basal growth, yet markedly enhanced radiation-induced loss of viability, caspase-3/7 activation, and Annexin V positivity across glioma, breast, prostate, and colorectal cancer models. This pattern is most consistent with XRRA1 acting as a stress-buffering factor, particularly during treatment, rather than as a constitutive driver of proliferation. In that sense, XRRA1 behaves like a therapy-emergent dependency: relatively dispensable under baseline conditions, but critical once cells are challenged by radiation damage ^[17]^. The newly added long-term survival and imaging data make that conclusion substantially stronger. XRRA1 loss disrupted spheroid architecture, reduced 2D colony foci, and sharply diminished clonogenic recovery as radiation dose escalated. Quantitative microscopy also showed that XRRA1 knockdown increased γH2AX even in the absence of irradiation and drove progressive γH2AX accumulation up to 8 Gy, implying that XRRA1 contributes not only to acute post-irradiation recovery but also to basal genome maintenance. The associated increase in DNA-PK puncta, the strong computational link to XRCC5, and the co-immunoprecipitation with Ku70 and XRCC4 place XRRA1 near the core non-homologous end-joining (NHEJ) machinery, where Ku and XRCC4-LIG4 coordinate broken-end alignment and repair. Although our study does not yet prove a direct enzymatic role for XRRA1 in end joining, the combined network, co-IP, and phenotype data strongly support its involvement in the containment or resolution of DNA double-strand break stress ^[3,30]^. This DNA-repair phenotype provides a plausible bridge to the innate immune findings. A growing body of work has shown that unresolved or misprocessed double-strand breaks can persist through mitosis, generate micronuclei, and expose self-DNA to cGAS, thereby initiating STING-TBK1-IRF3 signaling and interferon-stimulated gene expression. In our study, XRRA1 depletion plus radiation enhanced STING abundance, TBK1 and IRF3 phosphorylation, and a broad interferon-stimulated gene program, consistent with the idea that XRRA1 normally restrains the conversion of DNA damage into inflammatory signaling. Mechanistically, one parsimonious model is that XRRA1 helps channel radiation-induced lesions toward tolerable repair outcomes; when XRRA1 is lost, damaged DNA is more likely to persist, missegregate, or enter immune-sensing compartments, thereby amplifying cGAS–STING activation ^[6,7,9]^. Importantly, XRRA1 loss did not simply intensify cell death; it shifted the quality of cell death toward an immunogenic phenotype. Extracellular ATP release and cell-surface calreticulin exposure are among the best-established danger signals associated with immunogenic apoptosis, and both were markedly increased when XRRA1 silencing was combined with radiation. These findings, together with the interferon-stimulated gene response, support a model in which XRRA1 functions as a brake on radiation-induced immunogenic cell death. In practical terms, XRRA1 inhibition may therefore do more than radiosensitize tumor cells: it may also improve the immunologic visibility of dying irradiated cells, increasing the likelihood that radiotherapy engages productive tumor-immune crosstalk ^[13,24,31,32].^

The translational implications are therefore twofold. First, XRRA1 may serve as a biomarker. Its depletion in the CML proteome, favorable prognostic association, and induction in irradiated biopsies suggest that the meaning of XRRA1 depends on clinical context: baseline expression may reflect disease biology, whereas post-treatment expression may report adaptive radioresponse. Second, XRRA1 may be therapeutically actionable. Because its knockdown had modest effects under basal conditions but strong effects during irradiation, XRRA1 emerges as a plausible radiosensitization target. Any translational strategy will therefore need to define tumor selectivity, delivery strategy, and radiation dose/fractionation context carefully-especially given evidence that the immunogenic consequences of radiation depend strongly on how DNA damage is processed after treatment, which is shown in Figure 7 ^[5,6,9]^. XRRA1 is induced by radiation as part of an adaptive protective program; when present, it helps contain DNA damage and limit inflammatory conversion; when lost, DNA damage accumulates, apoptotic and clonogenic failure intensify, and cGAS-STING-driven immunogenic outputs emerge. In this framework, XRRA1 functions as a molecular checkpoint separating radiation tolerance from radiation-triggered immunogenic collapse. That model explains the major findings across all six result sections and identifies XRRA1 as a compelling candidate biomarker and therapeutic target for radiotherapy optimization.

**Figure 7:**
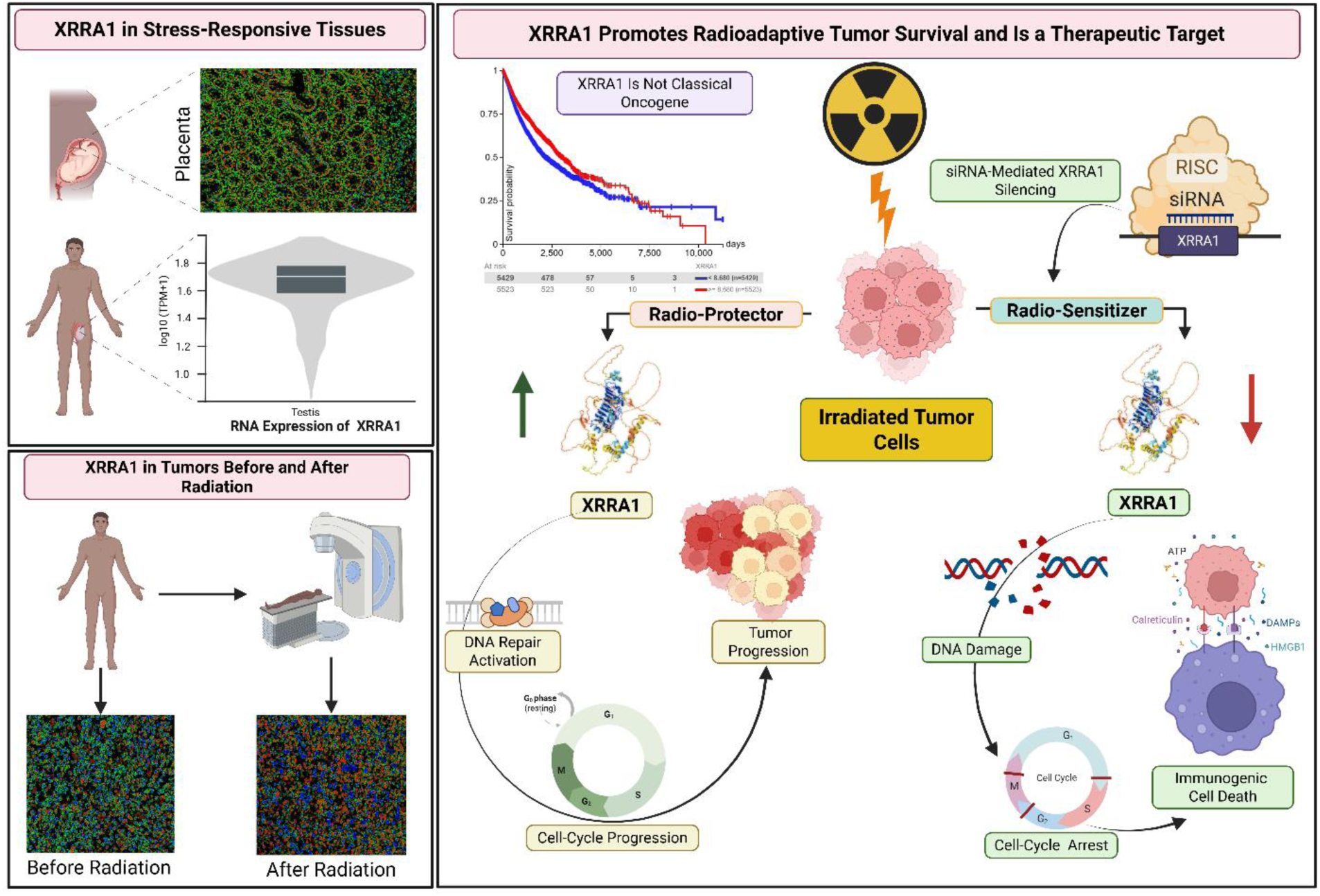
Graphical abstract shows a model of XRRA1-mediated radioadaptive tumor survival and radiosensitization. XRRA1 is expressed in stress-responsive tissues, including placenta and testis, and is increased in tumors after radiation exposure. In irradiated tumor cells, XRRA1 acts as a radioadaptive survival factor by supporting DNA damage tolerance, cell-cycle progression and tumor survival. By contrast, siRNA-mediated XRRA1 silencing functions as a radiosensitizing strategy, resulting in increased DNA damage, cell-cycle disruption, activation of cGAS-STING signaling, and induction of immunogenic cell death, accompanied by danger-associated outputs including ATP and calreticulin.

## Conclusions

Radiotherapy efficacy depends not only on the induction of DNA damage but also on whether damaged tumor cells adapt, die, or become immunogenic. We identify XRRA1 as a stress-adaptive regulator that buffers radiation-induced DNA damage signaling, limits cGAS-STING activation, and suppresses immunogenic cell death, thereby nominating XRRA1 as both a biomarker of radioadaptive stress and a candidate target for radiosensitization.

## CRediT authorship contribution statement

Tanvi Qamar: Investigation, Validation, Methodology, Formal analysis, Data curation. Saba Ubaid: Investigation, Methodology, Formal analysis. Vipin Kumar: Investigation, Validation, Methodology, Formal analysis, Resources. Mohammad Kashif: Investigation, Validation, Methodology, Formal analysis, Resources. Tanvi Singh: Data curation, Validation, Formal analysis, Writing-review & editing. Misba Majood: Data curation, Validation, Formal analysis. Ajay Kumar Singh: Investigation, Data curation, Formal analysis. Rashmi Kushwaha: Conceptualization, Supervision, Validation, Visualization, Resources, Methodology, Investigation, Formal analysis, Data curation. Vivek Singh: Conceptualization, Methodology, Investigation, Formal analysis, Data curation, Visualization, Software, Resources, Project administration, Funding acquisition, Supervision, Writing-original draft, Writing-review & editing.

## Ethics approval and consent to participate

The studies involving human participants were reviewed and approved by KGMU. Written informed consent to participate in this study was provided by the participants’ legal guardian/next of kin.

## Acknowledgements

English editing was carried out with ChatGPT5. After using this tool, the authors reviewed and edited the content as needed and took full responsibility for the content of the publication.

## Abbreviations

G/Gy: Gray
ANOVA: analysis of variance
ATP: adenosine triphosphate
CALR: calreticulin
cGAS: cyclic GMP–AMP synthase
CI: confidence interval
CML: chronic myeloid leukemia
DAB: 3,3′-diaminobenzidine
DAMPs: damage-associated molecular patterns
DeepLIIF: Deep Learning Inferred Immunofluorescence
DNA-PK: DNA-dependent protein kinase
DSB: DNA double-strand break
GEPIA3.0: Gene Expression Profiling Interactive Analysis 3.0
GTEx: Genotype-Tissue Expression
H-score: histological score
HMGA2: high mobility group AT-hook 2
HR: hazard ratio
ICD: immunogenic cell death
IFIT3: interferon-induced protein with tetratricopeptide repeats 3
IR: ionizing radiation
ISGs: interferon-stimulated genes
IHC: immunohistochemistry
KEA: kinase enrichment analysis
LC-MS/MS: liquid chromatography-tandem mass spectrometry
NHEJ: non-homologous end joining
OAS1: 2′-5′-oligoadenylate synthetase 1
OAS2: 2′-5′-oligoadenylate synthetase 2
PARP: poly(ADP-ribose) polymerase
PI: propidium iodide
siRNA: small interfering RNA
STING: stimulator of interferon genes
STRING: Search Tool for the Retrieval of Interacting Genes/Proteins
TBK1: TANK-binding kinase 1
TCGA: The Cancer Genome Atlas
TPM: transcripts per million
TBS: Tris-buffered saline
UPLC: ultra-performance liquid chromatography
XRCC5: X-ray repair cross-complementing protein 5
XRRA1: X-ray radiation resistance associated 1.

## Supplementary table legends

Supplementary Table 1: RT-PCR TaqMan probes list.

Supplementary Table 2: Antibodies Lists.

**Supplementary Table 3: LC–MS/MS protein identification metrics for healthy controls.** Complete LC–MS/MS protein identification dataset for peripheral-blood samples from healthy controls. Columns include protein annotation, molecular weight, isoelectric point, PLGS score, number of identified peptides, number of theoretical peptides, sequence coverage, precursor RMS mass error, number of product ions, digest peptides, modified peptides, product RMS mass error, and product RMS retention-time error. This table underlies the control proteome used for the overlap analysis in Fig. 1a.

**Supplementary Table 4: LC–MS/MS protein identification metrics for chronic myeloid leukemia samples.** Complete LC–MS/MS protein identification dataset for peripheral-blood samples from patients with chronic myeloid leukemia. Columns are as described for Supplementary Table 3. This table underlies the CML proteome used for the overlap analysis in Fig. 1a.

**Supplementary Table 5: STRING interaction dataset for the XRRA1-centred network.** STRING-derived interaction dataset used to generate the XRRA1-centred protein–protein interaction network shown in Fig. 1b. Fields include node 1, node 2, node 1 accession, node 2 accession, node 1 annotation, node 2 annotation, and STRING combined interaction score.

**Supplementary Table 6: Survival statistics for XRRA1 and XRRA1-associated genes.** Gene-wise survival analysis output used for Fig. 1c. Columns include gene symbol, Ensembl gene ID, regression coefficient, standard error, hazard ratio, 95% confidence interval, Z-score and *P* value for XRRA1 and genes within the XRRA1-associated interaction network.

**Supplementary Table 7: Top-ranked CRISPR codependencies associated with XRRA1.** DepMap CRISPR (Chronos) codependency dataset for XRRA1. Genes are ranked by correlation with XRRA1 dependency profiles. The table includes gene symbol, Entrez ID, dataset annotation and correlation coefficient and contains the positive and negative codependency genes used in Supplementary Fig. 1i.

**Supplementary Table 8: Top-ranked RNAi codependencies associated with XRRA1.** DepMap RNAi (DEMETER2) codependency dataset for XRRA1. Genes are ranked by correlation with XRRA1 dependency profiles. The table includes gene symbol, Entrez ID, dataset annotation and correlation coefficient and contains the positive and negative codependency genes used in Supplementary Fig. 1i.

**Supplementary Table 9: XRRA1 immunohistochemical ranking and clinicopathological annotation of the validation cohort.** Ranked summary of XRRA1 immunohistochemical staining across the clinical biopsy cohort. Pages 1–2 list sample rank, sample ID, IHC-positive fraction, and counts of weak (1+), moderate (2+) and strong (3+) cells. Pages 3–4 provide the corresponding H-score, radiation exposure status, radiation therapy cycle, age/sex, UHID, biopsy site, diagnosis, and representative original images. Pages 5–6 show the corresponding DeepLIIF-derived XRRA1, DAPI, hematoxylin, LAP2 and segmentation panels.

**Supplementary Table 10: DeepLIIF-derived digital H-score summary and scoring workflow for representative XRRA1-stained tissues.** Sheet 1 summarizes digital XRRA1 quantification for the representative tissues shown in Fig. 2, including radiation-exposed tissue, tissue without radiation history, placenta positive control, and negative control. Parameters include total nuclei analyzed, XRRA1-positive cells, positive-cell percentage, counts of weak (+1), moderate (+2) and strong (+3) cells, and the calculated H-score (0–300). Sheet 2 describes the digital scoring workflow, including segmentation source, positive-cell definition, intensity measurement from Marker.png, object filtering, pooled intensity thresholds for weak, moderate and strong staining, and the formula used for H-score calculation.

**Supplementary Table 11: Composition of the human XRRA1 ON-TARGET plus SMART pool siRNA reagent.** Composition of the human XRRA1 ON-TARGET plus SMART pool used for knockdown experiments. Columns include gene, species, product type, pool type, siRNA ID, catalogue number, target sequence, molecular weight, and extinction coefficient. The pool comprises four individual XRRA1-targeting siRNAs.

**Supplementary Table 12: Kinase enrichment analysis of XRRA1-associated proteins.** Ranked output of kinase enrichment analysis for XRRA1-associated genes or proteins. Columns include rank, kinase or gene set name, and Z-score. The table highlights the top predicted kinase associations used for interpretation of XRRA1-linked signalling networks.

**Supplementary Table 13: Quantitative microscopy workflow and per-nucleus analysis of XRRA1 and p(S139) γH2AX signals.** Quantitative image-analysis dataset underlying Supplementary Fig. 4e. Method summarizes the analysis pipeline, including dataset composition, treatment conditions, segmentation source, quantified channels, background subtraction method, positivity thresholds and control-based thresholding rule. Raw Nuclei contains per-nucleus measurements, including nuclear area, background-subtracted XRRA1 and γH2AX intensities, DAPI signal and binary positive calls. Field Summary reports per-field nuclei count, average signal intensities, positive fractions and background values. Condition Summary provides pooled condition-level averages, total nuclei analyzed, positive-cell fractions and fold changes relative to control. Segmentation documents the quality-control procedure used for nuclear segmentation. Because quantification was derived from composite display-normalized PNG panels rather than raw acquisition files, γH2AX trends should be interpreted as more robust than absolute XRRA1 red-channel measurements.

---

**Supplementary Figure 1:**
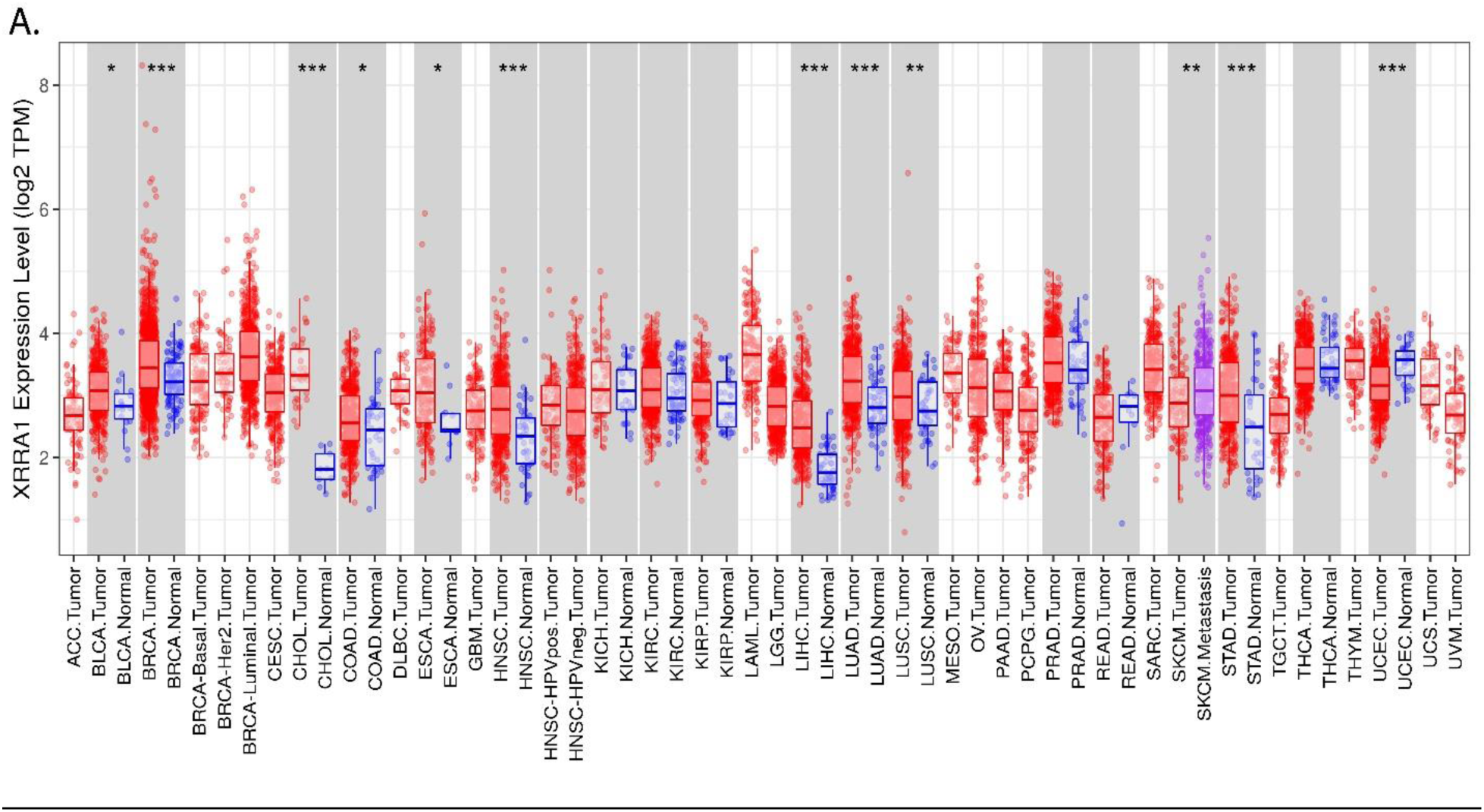
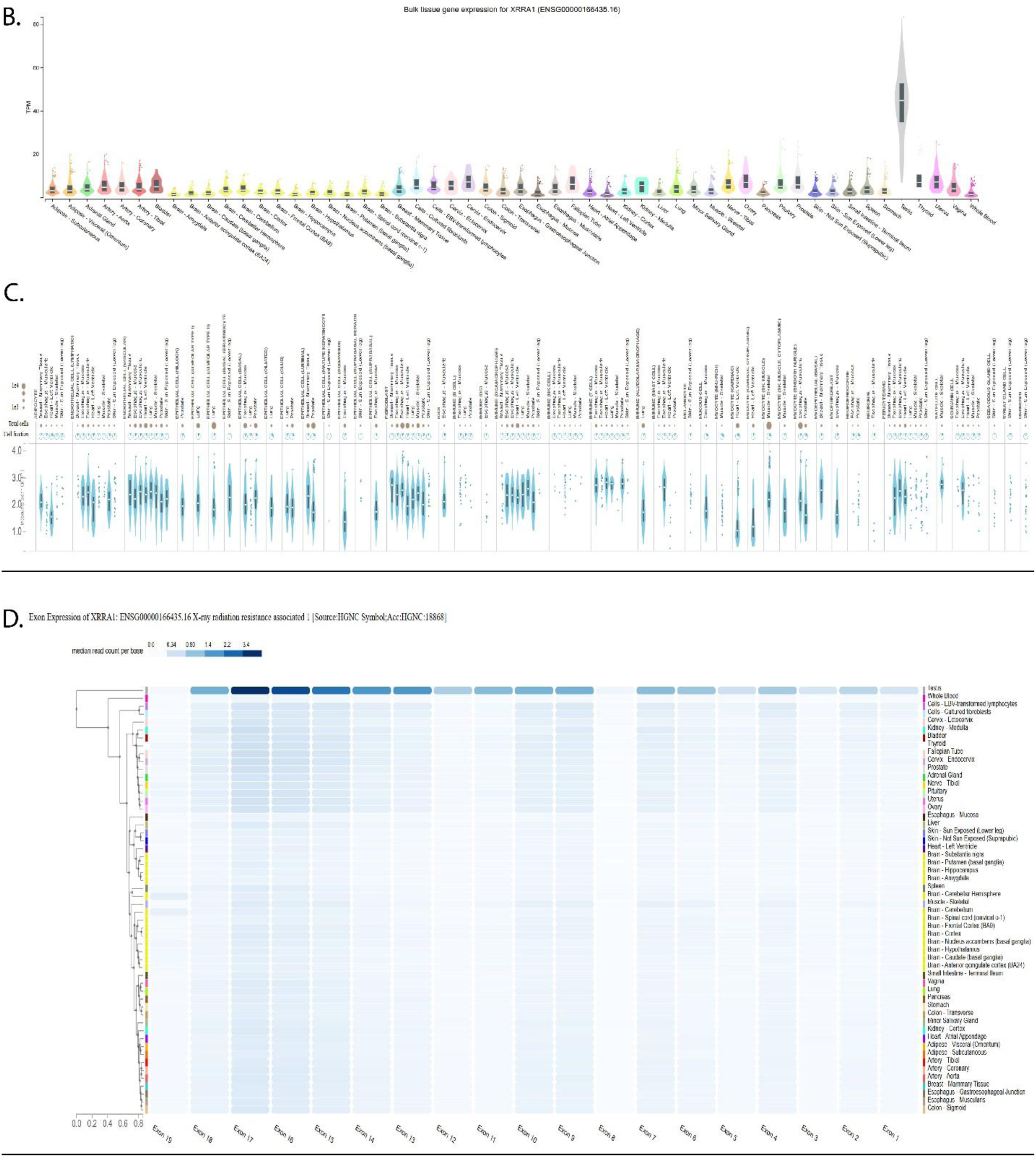
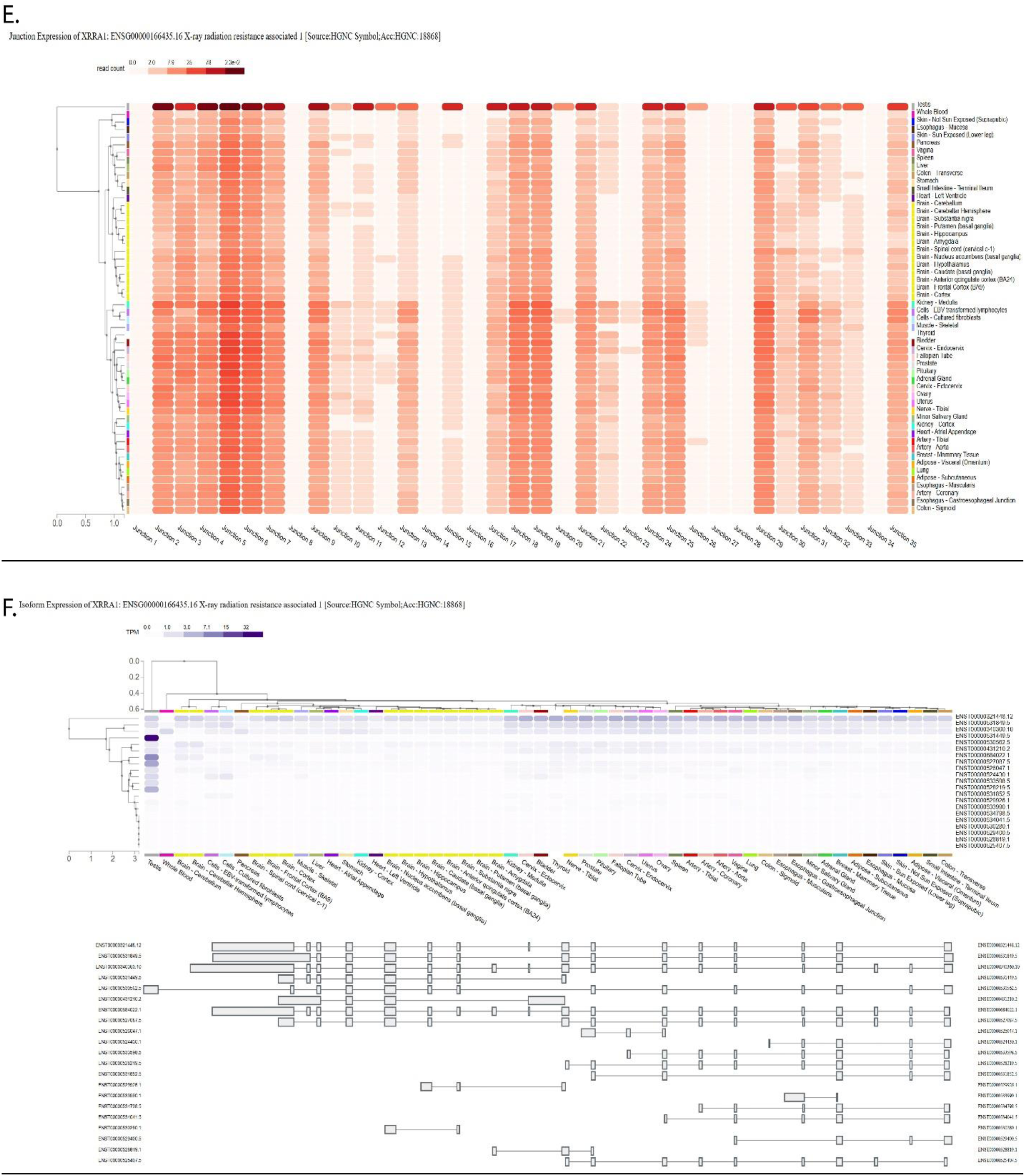
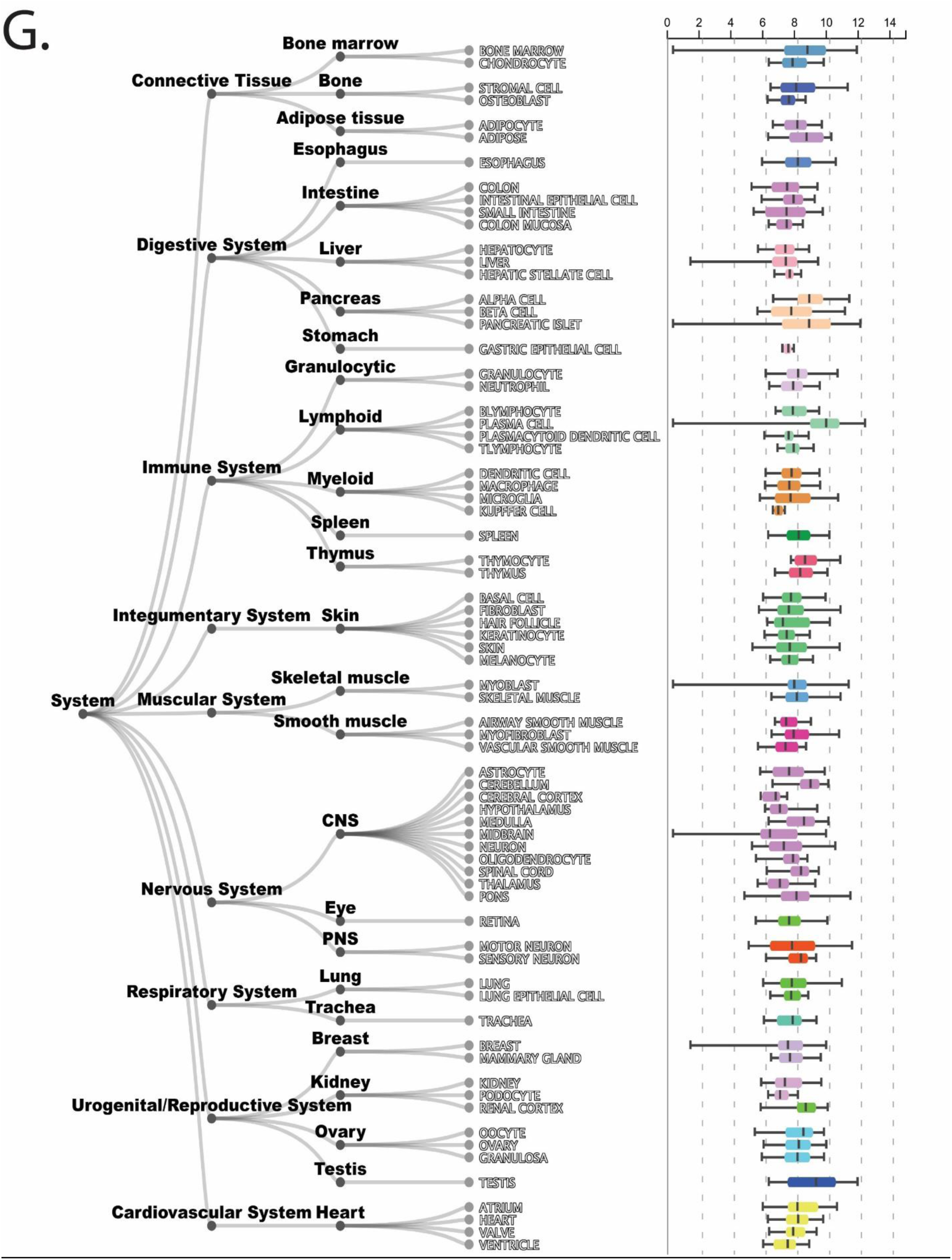
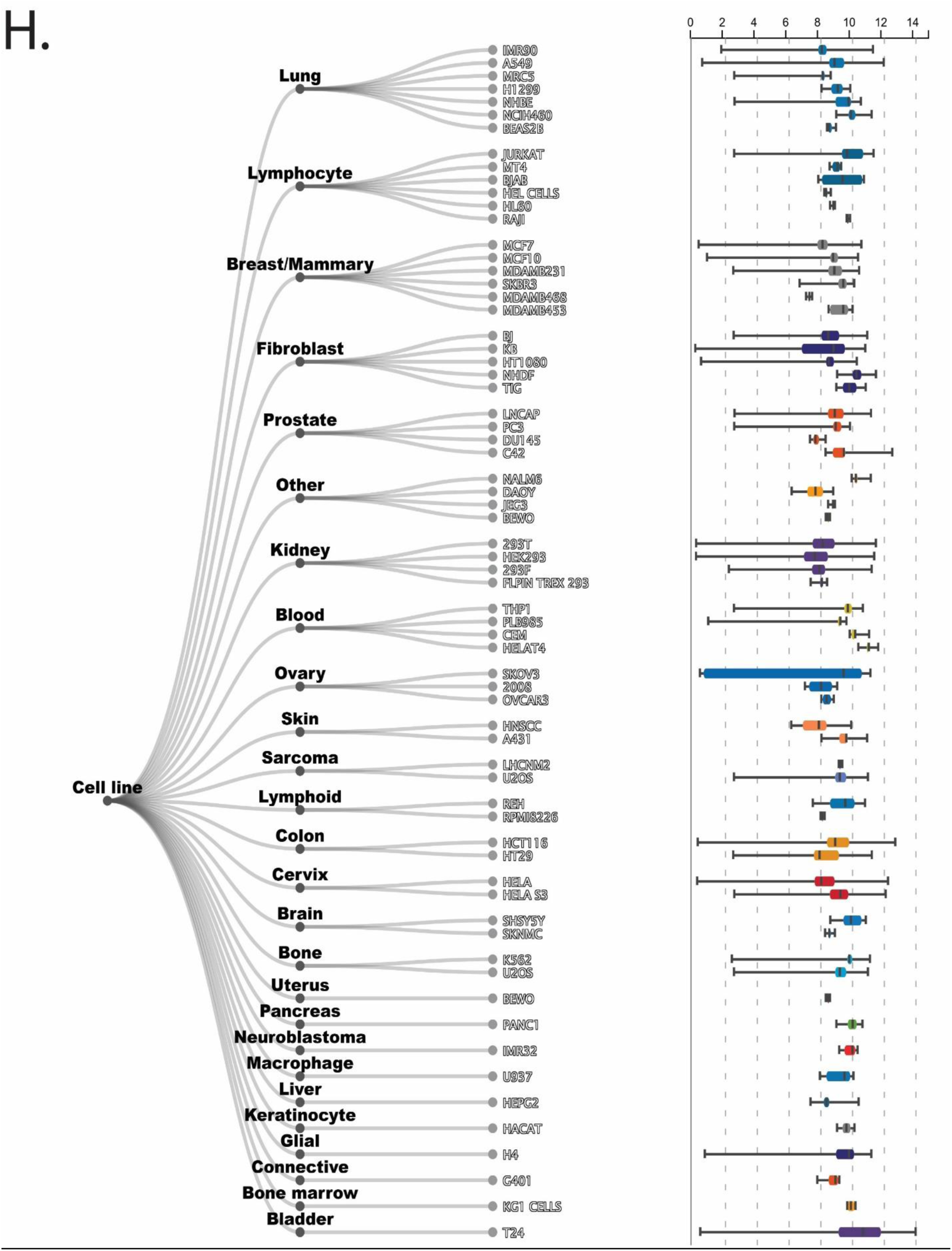
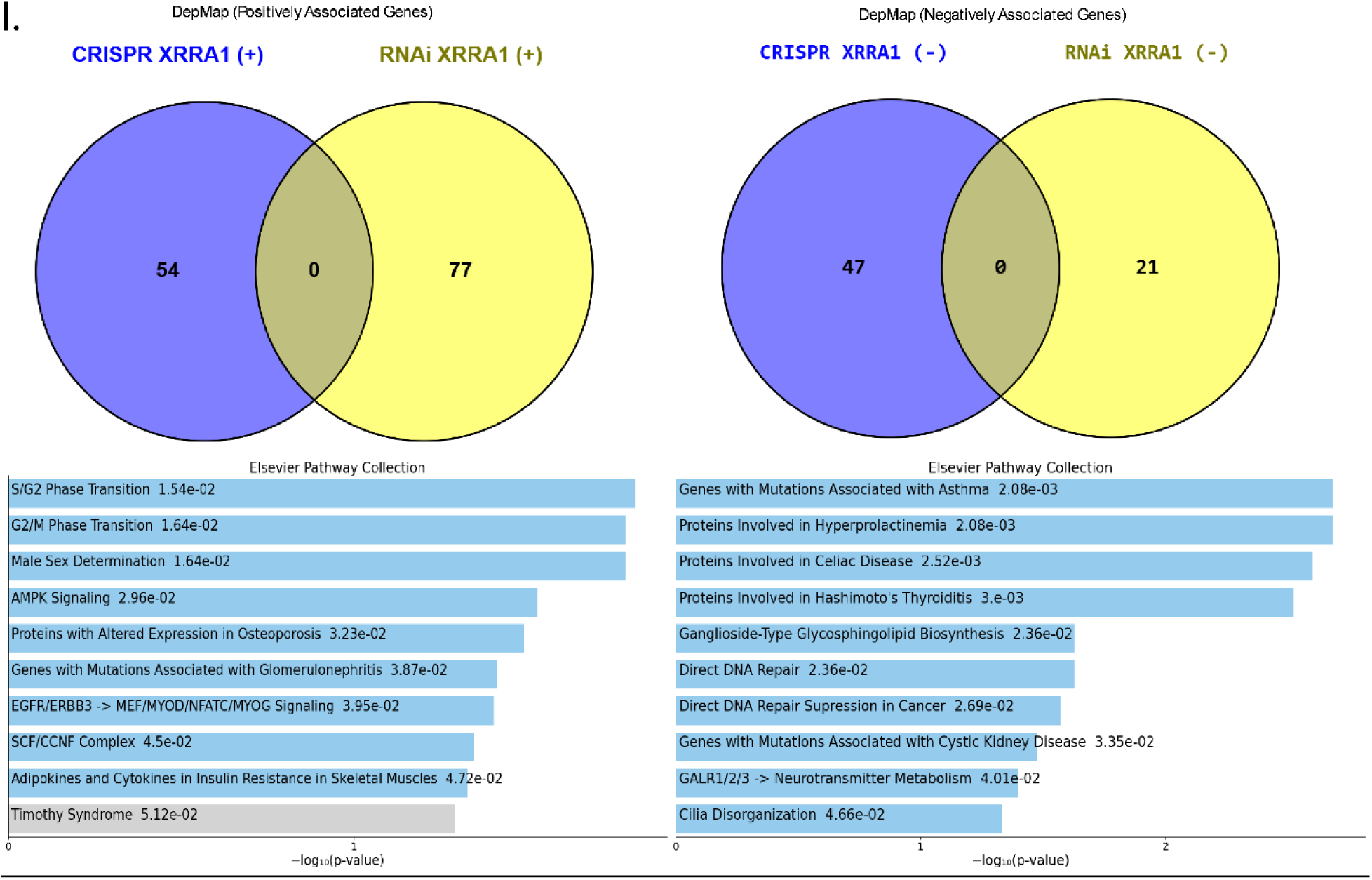
Expression landscape, transcript organization and dependency context of XRRA1. **S1A,** Pan-cancer comparison of XRRA1 mRNA expression across tumor and normal tissues. Box plots with overlaid data points show XRRA1 expression levels (log2 TPM) in tumor tissues (red) and normal tissues (blue); metastatic melanoma samples are shown in purple, where available. Cancer-type abbreviations follow standard TCGA nomenclature. Asterisks indicate statistically significant differences between groups. **S1B,** Bulk normal-tissue expression profile of XRRA1 across human tissues. Violin plots show TPM distributions across tissues, indicating broad basal expression with marked enrichment in selected tissues, including testis. **S1C,** Single-cell expression atlas of XRRA1 across human tissues. The upper tracks indicate the total number of cells analyzed and the fraction of XRRA1-expressing cells for each annotated cell type, whereas the lower violin plots show the distribution of XRRA1 expression at single-cell resolution. **S1D,** Exon-level expression heat map of XRRA1 across normal tissues. Columns correspond to XRRA1 exons 1-19, and color intensity represents median read count per base. Tissues are hierarchically clustered according to exon-level expression patterns. **S1E,** Junction-level expression heat map of XRRA1 across normal tissues. Columns correspond to splice junctions 1–35, and color intensity represents junction read count. Tissues are hierarchically clustered according to splice-junction usage. **S1F,** Isoform expression landscape of XRRA1 across normal tissues. The upper heat map shows clustered isoform-level expression (TPM) across tissues, and the lower panel shows the exon–intron architecture of annotated XRRA1 transcripts. **S1G,** System-level normal-tissue atlas of XRRA1. The branching diagram organizes XRRA1 expressions across major body systems, tissues and representative cell types, including bone marrow, immune, digestive, integumentary, nervous, respiratory, urogenital/reproductive and cardiovascular compartments. Box plots at right summarize relative XRRA1 expression across the corresponding tissue and cell-type categories. **S1H,** Cell-line expression atlas of XRRA1. The branching diagram groups cultured cell lines by tissue or lineage of origin, and the accompanying box plots summarize relative XRRA1 expression across the indicated categories. **S1I,** DepMap codependency analyses for XRRA1. Venn diagrams compare positively associated XRRA1 gene sets identified by CRISPR and RNAi screens (54 and 77 genes, respectively) and negatively associated gene sets (47 and 21 genes, respectively). Bar plots show Elsevier Pathway Collection enrichment for the corresponding positive and negative gene sets; bar length indicates −log10(*P*). Gene-level correlation data are provided in Supplementary Tables 7 and 8.

**Supplementary Figure 2:**
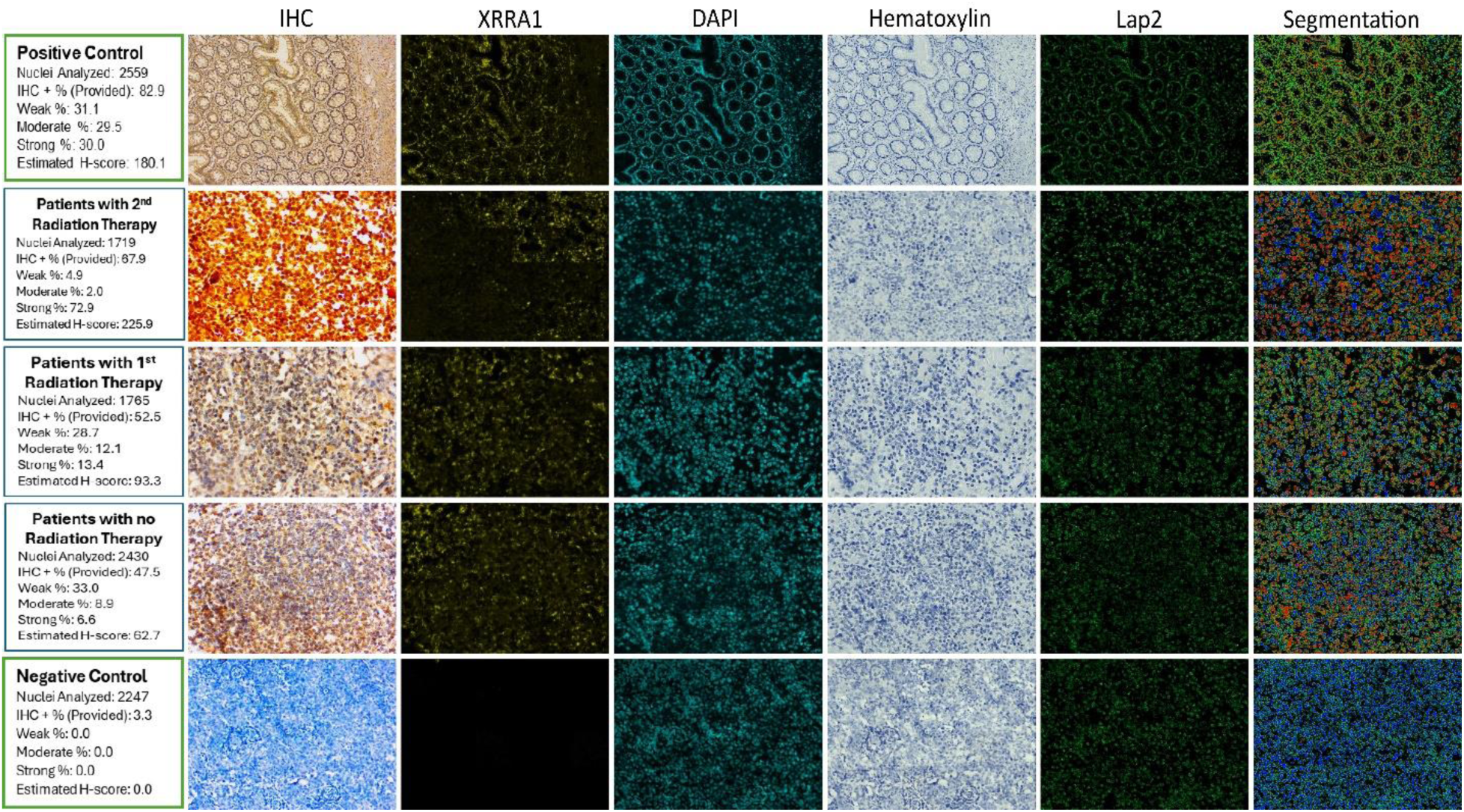
Extended digital pathology analysis of XRRA1 across radiation exposure groups. Representative XRRA1 immunohistochemistry and DeepLIIF-derived channel decomposition for positive control, second radiation therapy, first radiation therapy, no radiation therapy, and negative control groups. For each condition, the IHC, XRRA1, DAPI, hematoxylin, LAP2, and segmentation panels are shown. Annotation boxes summarize the number of nuclei analyzed, the percentage of IHC-positive cells, the proportions of weak, moderate and strong staining, and the estimated H-score for each representative field.

**Supplementary Figure 3:**
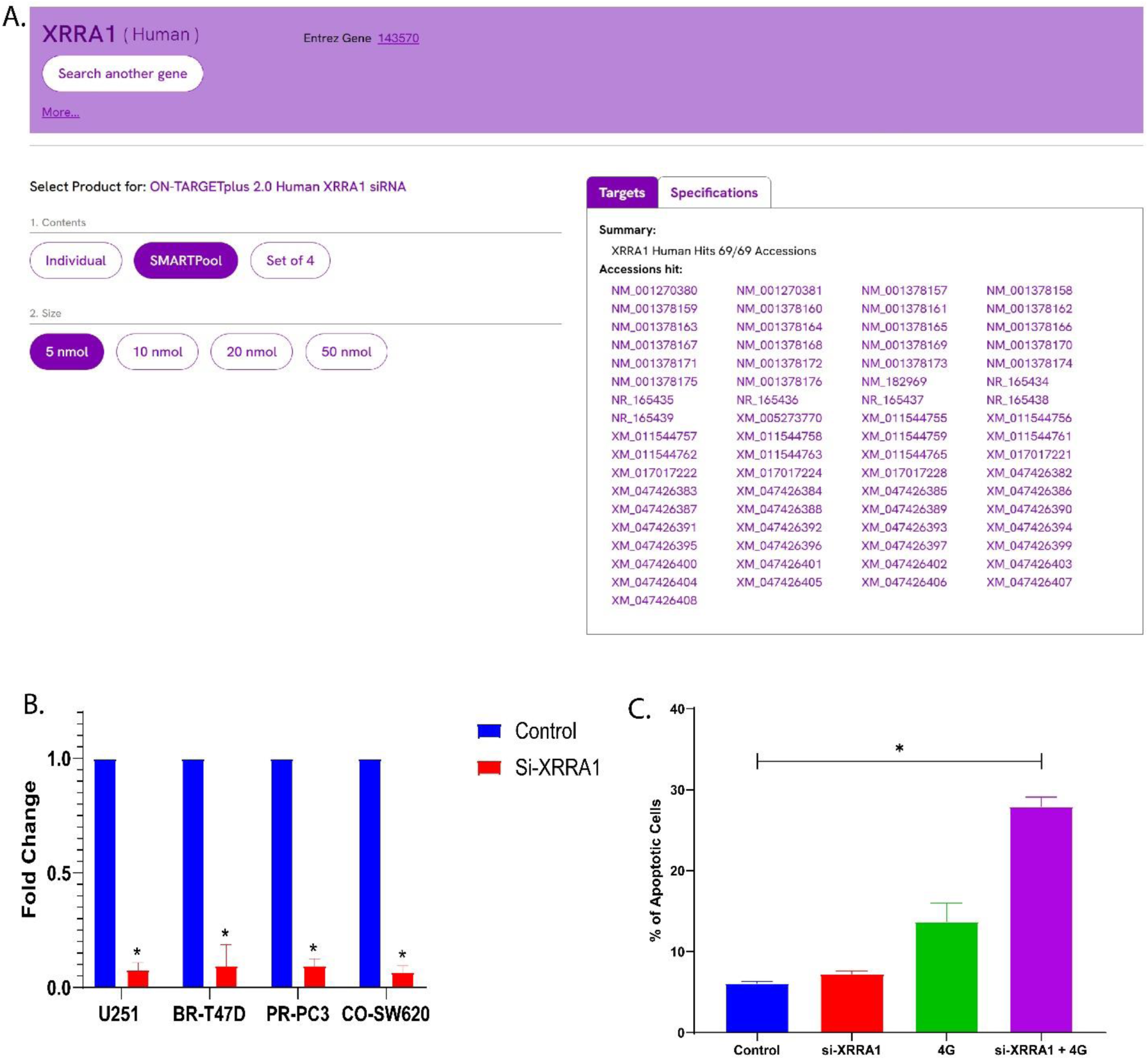
Validation of XRRA1-targeting siRNA and quantification of apoptosis following XRRA1 depletion and irradiation. **S3A,** Transcript coverage summary for the ON-TARGET plus 2.0 Human XRRA1 siRNA SMART pool, showing predicted targeting of XRRA1 accessions. **S3B,** Validation of XRRA1 knockdown efficiency in U251, BR-T47D, PR-PC3 and CO-SW620 cells. XRRA1 expression is shown as fold change relative to control, with control normalized to 1. **S3C,** Quantification of apoptotic cells under control, si-XRRA1, 4 Gy, and si-XRRA1 + 4 Gy conditions. Combined XRRA1 silencing and irradiation produced the greatest apoptotic response. Error bars indicate variation among replicate measurements. Statistical significance is indicated in the plot.

**Supplementary Figure 4:**
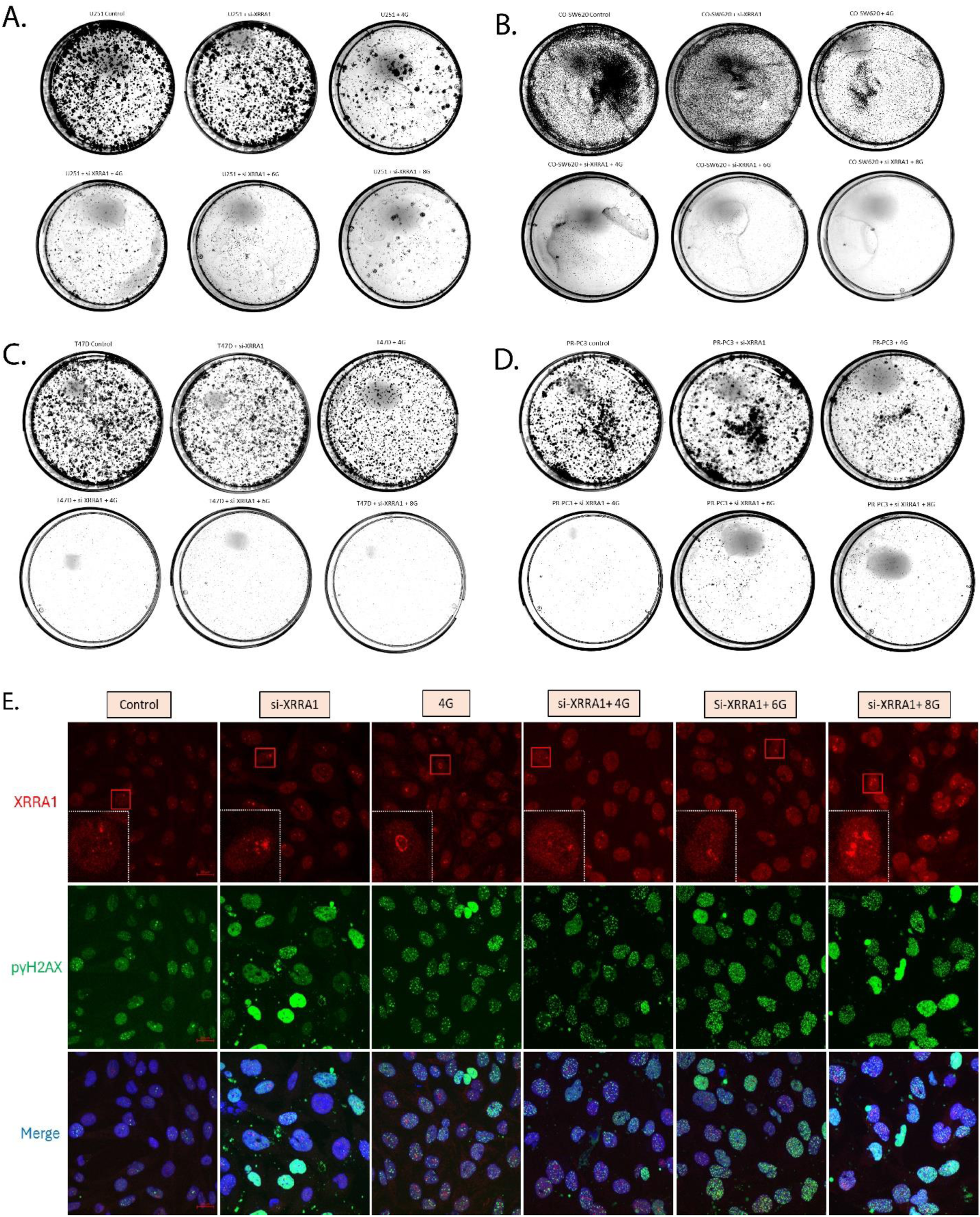
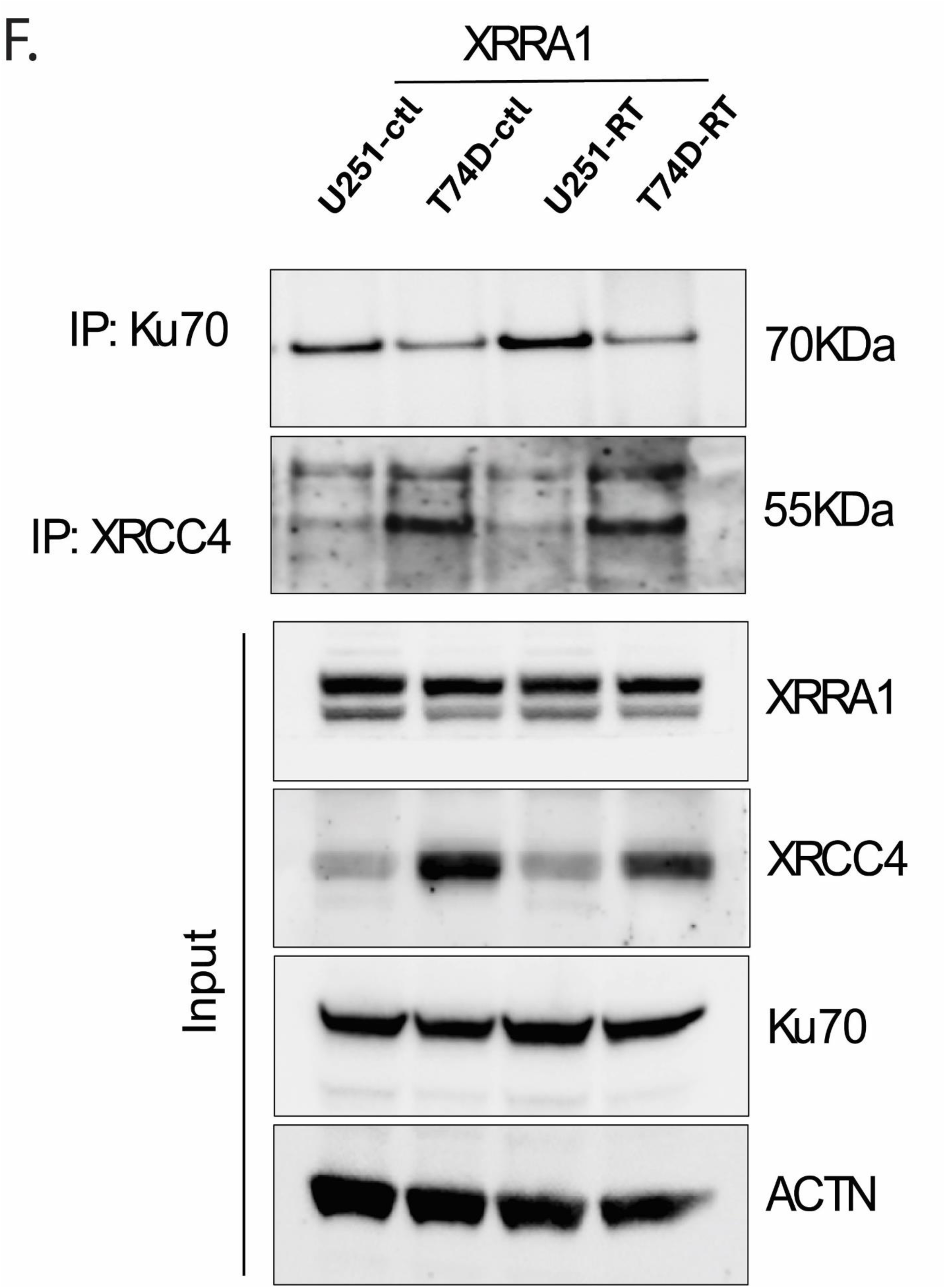
XRRA1 loss reduces clonogenic survival and augments dose-dependent DNA damage signaling. **S4A-S4D,** Representative clonogenic plates from U251 (a), CO-SW620 (b), T47D (c) and PR-PC3 (d) cells under control, si-XRRA1, 4 Gy, si-XRRA1 + 4 Gy, si-XRRA1 + 6 Gy, and si-XRRA1 + 8 Gy conditions. Across all four cell lines, XRRA1 silencing enhanced the inhibitory effect of radiation on clonogenic outgrowth, particularly at higher doses. **S4E,** Dose-escalation immunofluorescence images showing XRRA1 (red), p(S139) γH2AX (green), and merged images with nuclear counterstain (blue) under control, si-XRRA1, 4 Gy, si-XRRA1 + 4 Gy, si-XRRA1 + 6 Gy, and si-XRRA1 + 8 Gy conditions. Boxed regions indicate magnified areas. Quantitative analysis is provided in Supplementary Table 13. **S4F,** Co-immunoprecipitation and input immunoblot analysis in U251 and T47D cells under control and irradiated conditions. Upper panels show Ku70 and XRCC4 immunoprecipitates; lower panels show the corresponding input lysates immunoblotted for XRRA1, XRCC4, Ku70 and ACTN.

